# Characterizing Genetic Circuit Components in *E. coli* towards a *Campylobacter jejuni* Biosensor

**DOI:** 10.1101/290155

**Authors:** Natalia Brzozowska, Jane Gourlay, Ailish O’Sullivan, Frazer Buchanan, Ross Hannah, Alison Stewart, Hannah Taylor, Reuben Docea, Greig McLay, Ambra Giuliano, James Provan, Katherine Baker, Jumai Abioye, Julien Reboud, Sean Colloms

## Abstract

*Campylobacter jejuni* is responsible for most cases of bacterial gastroenteritis (food poisoning) in the United Kingdom. The most common routes of transmission are by contact with raw poultry. Current detection systems for the pathogen are time-consuming, expensive or inaccessible for everyday users. In this article we propose a cheaper and faster system for detection of *C. jejuni* using a synthetic biology approach. We aimed to detect *C. jejuni* by the presence of xylulose, an uncommon bacterial capsular saccharide. We characterized two sugar-based regulatory systems that displayed potential to act as tools for detection of xylulose. Using a two-plasmid reporter system in *Escherichia coli*, we investigated the regulatory protein component (MtlR) of the mannitol operon from *Pseudomonas fluorescens*. Our findings suggest that the promoter of *mtlE* is activated by MtlR in the presence of a variety of sugar inducer molecules, and may exhibit cross-activity with a native regulator of *E. coli*. Additionally, we engineered the L-arabinose transcriptional activator (AraC) of *E. coli* for altered ligand specificity. We performed site-specific saturation mutagenesis to generate AraC variants with altered effector specificity, with an aim to generate a mutant activated by xylulose. We characterized several mutant AraC variants which have lost the ability to respond specifically to the native L-arabinose effector. We promote this technique as a powerful tool for future iGEM teams to create regulatory circuits activated by novel small molecule ligands.

## Introduction

*Campylobacter jejuni* is a Gram-negative, microaerophilic, corkscrew-shaped bacteria which has been implicated as being one of the most common causes of human gastroenteritis worldwide [1][2][3]. Infection with *C. jejuni* causes common symptoms such as diarrhoea, abdominal pain, fever, headache, nausea and vomiting [3]. *C. jejuni* is harboured by poultry, though it has been reported in other meat products, raw milk, and in untreated drinking water [3]. The high prevalence of *C. jejuni* makes it an interesting target for synthetic biology-based solutions.

Traditional methods for detection of *C. jejuni* include culture-based techniques, which are relatively cheap to perform and require less training than other methods [4]. However, they are incredibly time and labour intensive and therefore, in recent years, a move has been made towards use of rapid detection testing [5]. Such techniques include enzyme immunoassay and lateral flow systems, which require only one to two hours to give a result [6]. However, use of these methods requires highly trained employees, and so detection of *C. jejuni* in an industrial or agricultural setting would require outsourcing to specialists. In addition, a comparison of three rapid detection systems demonstrated a high number of false negative results, which is a drawback when considering detection of *C. jejuni* to reduce the incidence of disease outbreaks [7].

To reduce incidences of food poisoning by *C. jejuni*, and to improve upon current methods of detection, we decided to create a biosensor that was able to detect the presence of *C. jejuni* quickly and accurately. We envisaged a two-part biosensor that would specifically require two sensory inputs associated with *C. jejuni* to report a reliable result. The first molecule we identified as a marker for *C. jejuni* was autoinducer-2 (AI-2) [9]. AI-2 is a secreted quorum-sensing molecule. However, many varied gram-positive and gram-negative bacterial species sense their population density and surrounding bacterial environment using this molecule [10]. On the other hand, this ubiquity meant that AI-2 gene regulation was well characterised, with prior iGEM teams having worked on the *E. coli* AI-2 quorum sensing regulatory system [11][12]. Therefore, we searched for another marker with greater specificity to *C. jejuni* to use in tandem with AI-2.

Xylulose is a rare sugar found incorporated within the polysaccharide capsule of *C. jejuni* [8]. The presence of xylulose is uncommon in bacterial polysaccharide capsules [8]. Additionally, the glyosidic bonds which incorporate xylulose were found to be extremely acid-labile [8], providing a possibility to release the molecule from the capsule and allow for whole-cell based detection. For the detection of xylulose two possible avenues were explored. One exploits the mannitol metabolism operon of *Pseudomonas fluorescens* [13]. It has been previously reported that xylulose acts as a direct inducer of the regulatory protein MtlR, activating transcription from the p_*mtlE*_ promoter [14]. To investigate its suitability for biosensor, we characterized the p_*mtlE*_/MtlR regulatory system in *E. coli*.

Use of a gene regulatory system outside the context of the native organism may be problematic. For this reason, we aimed to construct an alternative xylulose sensor, by exploiting components of the L-arabinose operon, native to *E. coli* [15]. Several previous studies have shown that its regulatory protein, AraC, can be engineered to activate transcription in response to non-native small molecules [16][17][18]. Site-saturation mutagenesis of residues positioned within the ligand-binding pocket of AraC (Fig 1), coupled with fluorescence-based cell sorting, allowed isolation of AraC variants with altered effector specificity [16]. Based on these findings we aimed to use multiple site-saturation mutagenesis and fluorescence-based screening to generate mutant AraC responsive specifically to xylulose.

**Fig 1.**
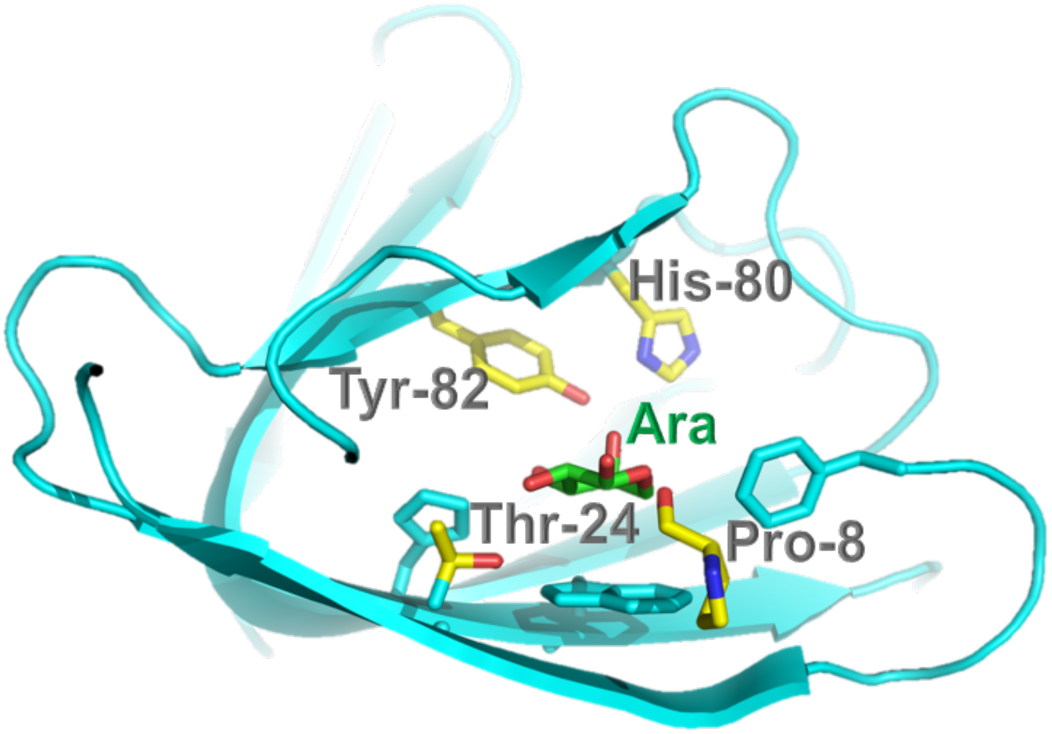
Crystal structure of AraC biding pocket with bound L-arabinose (Ara). The four key residues which are important for ligand binding pocket are indicated. Structure was generated using PyMOL.

## Materials and Methods

### Standard molecular biology techniques

All protocols used during this work, including a standardised set for routine laboratory techniques, are detailed in S3 File. All ligation reactions described herein were transformed into DH5α commercial chemically competent *E. coli* cells, plated on Lagar containing the required antibiotics, and grown at 37°C overnight. All plasmid constructs herein were verified by diagnostic restriction digest and DNA sequencing prior to use or further subcloning.

### Plasmid design for expression of p_*mtlE*_/MtlR regulatory system

All plasmids were constructed using BioBrick Standard Assembly. The biobrick p_*tet*_ promoter (R0040) followed by a medium-strength ribosome binding site (RBS) B0032 was ordered for synthesis by Integrated DNA Technologies (IDT) as complementary oligos, that were annealed and ligated into a pSB1C3 plasmid backbone. *mtlR* coding sequence was ordered as a gBlock Gene Fragment by IDT and inserted behind the R0040+B0032 promoter/RBS part in pSB1C3. This produced the regulatory plasmid BBa_K2442202 (Fig 2).

**Fig 2.**
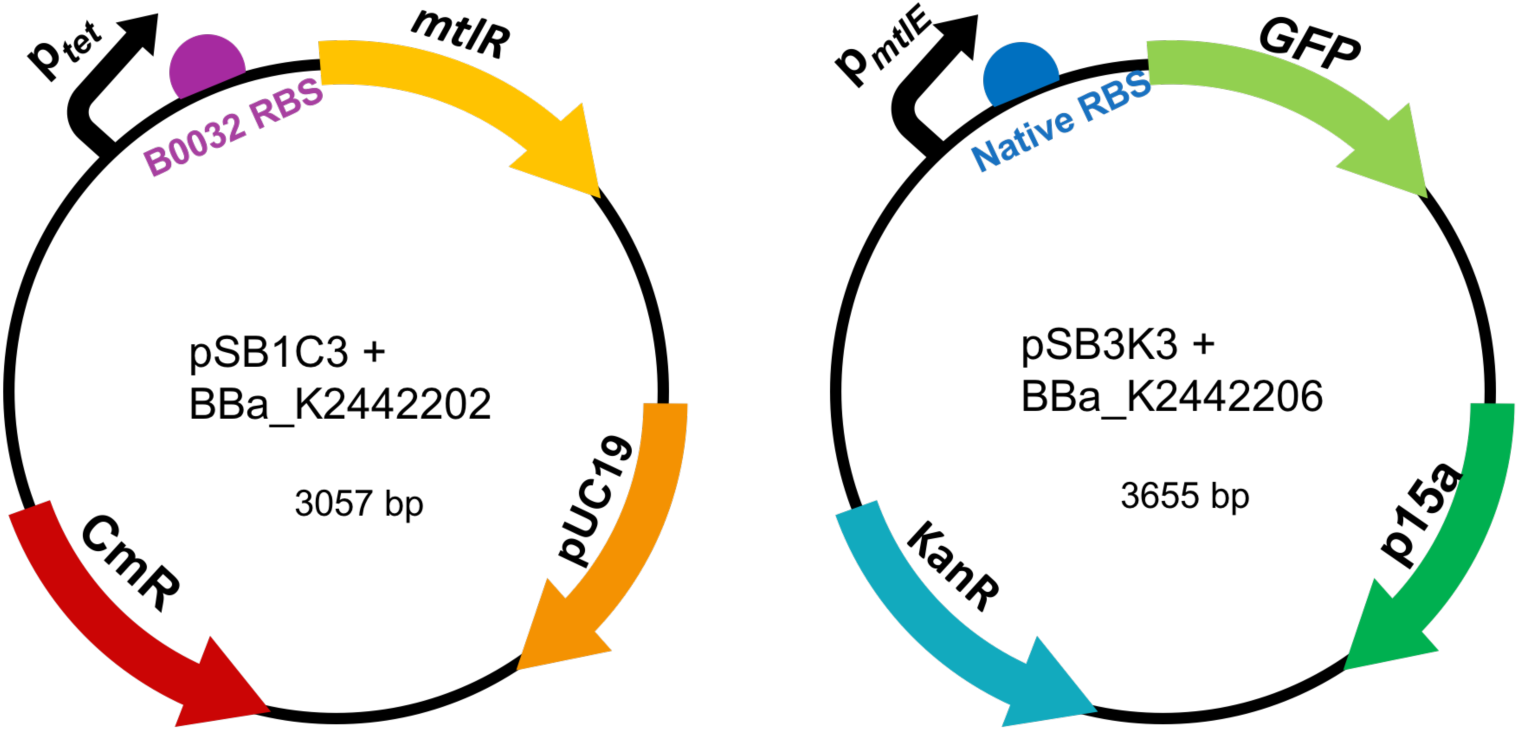
Diagrams of the regulatory plasmid (left) and GFP reporter plasmid (right) for the expression of the MtlR/p_mtlE_ regulatory system.

The sequence of the p_*mtlE*_ promoter with its native RBS was obtained from Liu *et al*. (2015) [14] and supplied as oligo by IDT. A reporter plasmid was constructed by ligating GFP coding sequence (BBa_E0040; obtained from iGEM Distribution Kit) downstream of the p_*mtlE*_ promoter into pSB1C3 backbone. The resulting plasmid BBa_K2442206 is shown in Fig 2.

### Plasmid design for expression of p_*BAD*_/AraC regulatory system

Fragments containing parts R0011 (LacI-regulated promoter) upstream of ribosome-binding site (RBS) B0032 were synthesised by IDT as oligonucleotides. Wild type *araC* sequence as described by Miyada *et al*. (1980) [19] was amplified from BBa_I0500 using primers *araC_BBPre_F* and *araC_BBSuf_R* (Table 1) to introduce BioBrick prefix and suffix to both ends of WT *araC*. The PCR product was subsequently inserted downstream of R0011 promoter and B0032 RBS in pSB1C3 backbone to generate the regulatory plasmid BB_K2442104 (Fig 3).

**Fig 3.**
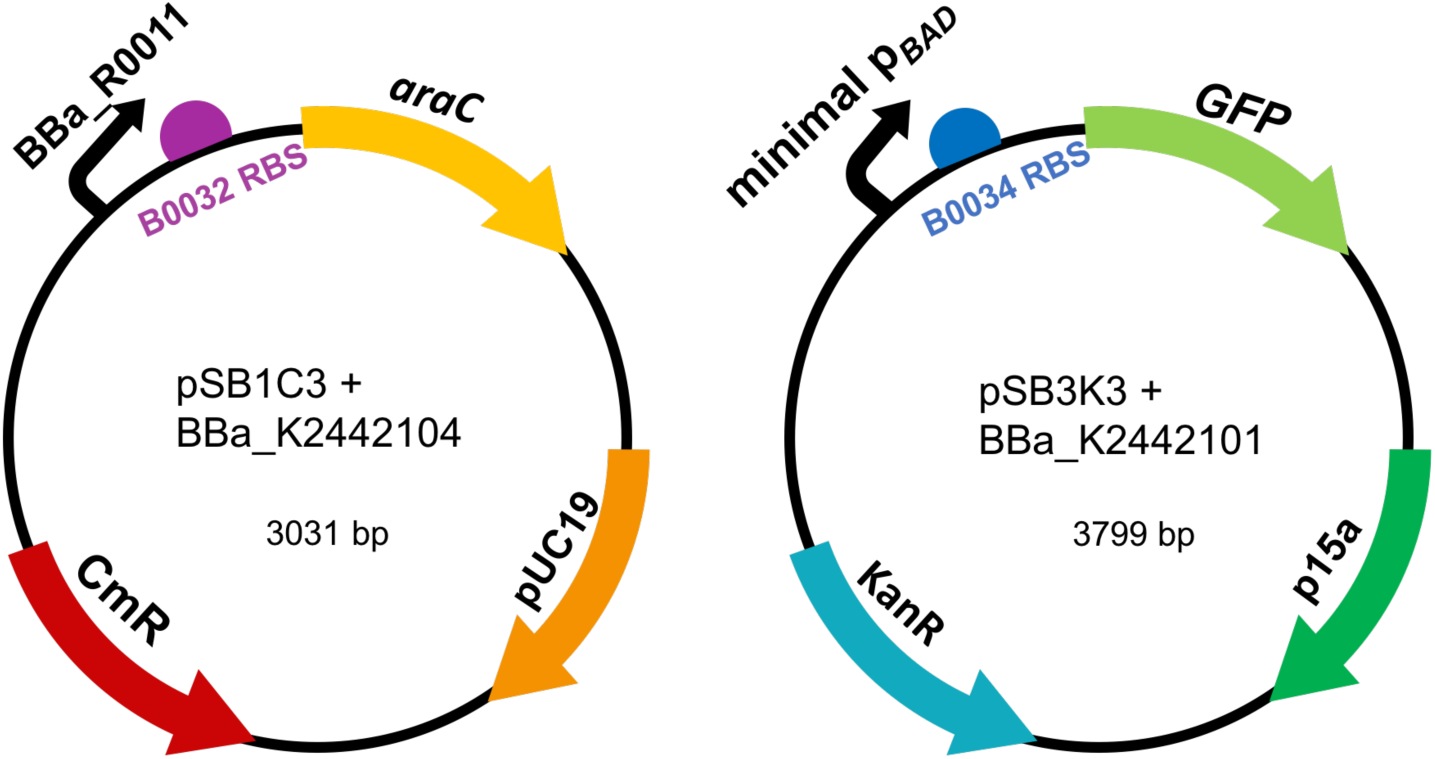
Diagrams of the regulatory plasmid (left) and GFP reporter plasmid (right) for the expression of the AraC/p_BAD_ regulatory system.

Minimal p_*BAD*_ promoter was synthesised by IDT as a gBlock (see S1 Fig for full sequence). The fragment was ligated into pSB3k3, upstream of part BBa_I13500 (containing B0034 RBS and GFP). This resulted in the final reporter plasmid BBa_K2442102 (Fig 3).

### Mutant AraC Library Construction

Site sirected mutagenesis by PCR was performed according to the protocol in S3 File. QIAQuick PCR purification of the product was performed according to the Qiagen protocol (S3 File). The purified PCR product was ligated into pSB1C3 backbone, downstream of R0011 promoter and B0032 RBS. Ligation reaction was ethanol-precipitated, and transformed into electrocompetent DH5α by electroporation. This resulted in the *araC* mutant library. Colonies carrying the mutant library were washed off transformation plates with 1.5ml double distilled H_2_O, (ddH_2_O) then transferred to a 1.5ml tube. The cell pellet was centrifuged at room temperature, the supernatant was then discarded and pellet resuspended in 1.5ml ddH_2_O. This centrifugation-resuspension step was repeated 3 times to remove agar plate debris. Plasmid DNA was extracted using the Qiagen Plasmid MiniPrep Kit with 2 of each PB buffer and PE buffer wash steps to ensure maximum extract purity. Sequencing primer VF2 was then added and sample sent for sequencing.

### Characterization of the p_*mtlE*_/MtlR system activity in *E. coli*

To characterize activity of the p_*mtlE*_/MtlR in *E. coli*, we studied the levels of GFP fluorescence using a 96 well plate in the FLUOstar Omega plate reader (BMG Labtech). Each strain was grown to saturation in an overnight culture of LB with appropriate antibiotics for the relevant plasmids (chloramphenicol and kanamycin). Then the culture was diluted 1:100 with fresh LB+antibiotic and placed into a black bottomed 96 well plate. All readings were performed in the plate reader at 37°C shaking at 200 RPM. GFP fluorescence was measured (excitation at 485nm and emission at 530nm) every hour for 8 hours.

### Mutant Screening

Plasmid DNA containing the *araC* mutant libraries were transformed into DS941 strain *E. coli* carrying the reporter plasmid K2442102 (in pSB3K3 vector). Transformants were plated on LB agar medium containing chloramphenicol plus kanamycin (to select for K2442102 and K2442104) plus one of the tested inducers: arabinose, xylose or decanal. Concentrations were 40mM for arabinose and xylose and 2mM for decanal. As a control test, transformants were also plated on LB medium containing chloramphenicol and kanamycin only. Fluorescence images of the conditional transformation plates were obtained using a GE-Healthcare Typhoon FLA-9500 laser scanner. Excitation was recorded at 473nm, emission was recorded using 520-540nm filter. Colonies which exhibited fluorescence were observed as dark colonies on the scan. Out of those which appeared dark on xylose or decanal plates, 200 were replica short-streaked onto plates containing xylose, arabinose, decanal, or no additive. Each colony of interest was picked and then immediately short-streaked onto each new condition plate using the same toothpick, into the same position using grids. This method of replica plating ensures the plates can later be aligned and short streaks of the same origin directly compared between the plates. After overnight incubation at 37°C, fluorescence scans of each plate were again obtained using the laser scanner.

### Liquid culture fluorescence assay for AraC mutants

Colonies of interest were inoculated into L-broth containing chloramphenicol and kanamycin. Each liquid culture was grown overnight at 37°C, shaking at 225 rpm. The following day the culture was diluted 1:100 into fresh L-broth containing the above antibiotics plus one of the following inducer conditions:

- 40mM xylose
- 40mM arabinose
- 2mM decanal

200µl of each culture to be tested was placed in a well of a clear-based 6-well plate, then incubated at 37°C shaking at 300 rpm for 12 hours in a BMG FLUOstar Omega fluorescence plate reader. GFP fluorescence of each culture was measured at 1 hour time intervals for the duration of the culture experiment, using excitation wavelength 485nm and emission wavelength 530nm. Cell growth was simultaneously tracked by measuring optical density at 600nm.

## Results

### P_*mtlE*_ is regulated by MtlR and other native regulatory proteins in *E. coli*

From the literature, we identified a regulatory system that responds to xylulose-the mannitol-inducible promoter from *P. fluorescens* and its regulatory protein MtlR [14]. To utilize these parts in our dual-input biosensor, their activity needed to be characterized in *E. coli*. Two constructs were assembled: the regulatory plasmid with a constitutively active p_*tet*_ promoter driving expression of the MtlR protein, and the reporter plasmid containing p_*mtlE*_ promoter regulating expression of GFP (Fig 1).

To test whether MtlR can activate p_*mtlE*_ in *E. coli*, we measured GFP fluorescence in *E. coli* carrying the reporter plasmid alone, or both the regulatory and reporter plasmids. The experiment was done in presence of 6 structurally similar sugars (ribose, fructose, xylose, mannitol, arabinose and sorbitol) [14], to investigate substrate specificity of MtlR. Xylulose itself was not tested due to budgetary constraints. Fluorescence levels were compared to basal fluorescence levels of DH5α cells, not expressing either plasmid. Cells expressing the reporter plasmid alone showed higher levels of fluorescence than empty cells, in presence of all sugars tested (Fig 4 and S1 File). This suggests that P_*mtlE*_ may interact with native *E. coli* proteins. GFP fluorescence levels further increased in cells expressing both MtlR and the reporter plasmid (Fig 4 and S1 File). This supports previous findings that MtlR functions as an activator of p_*mtlE*_ in *E. coli*. Although we were unable to test xylulose, we found that p_*mtlE*_ was induced in presence of a number of structurally similar sugars, showing highest response to ribose and sorbitol (Fig 4 and S1 File). We conclude that p_*mtlE*_ promoter functions in *E. coli*, and MtlR acts as its activator. However, the *P. fluorescens* p_*mtlE*_ promoter is not strictly regulated by MtlR when expressed in *E. coli*.

**Figure 4.**
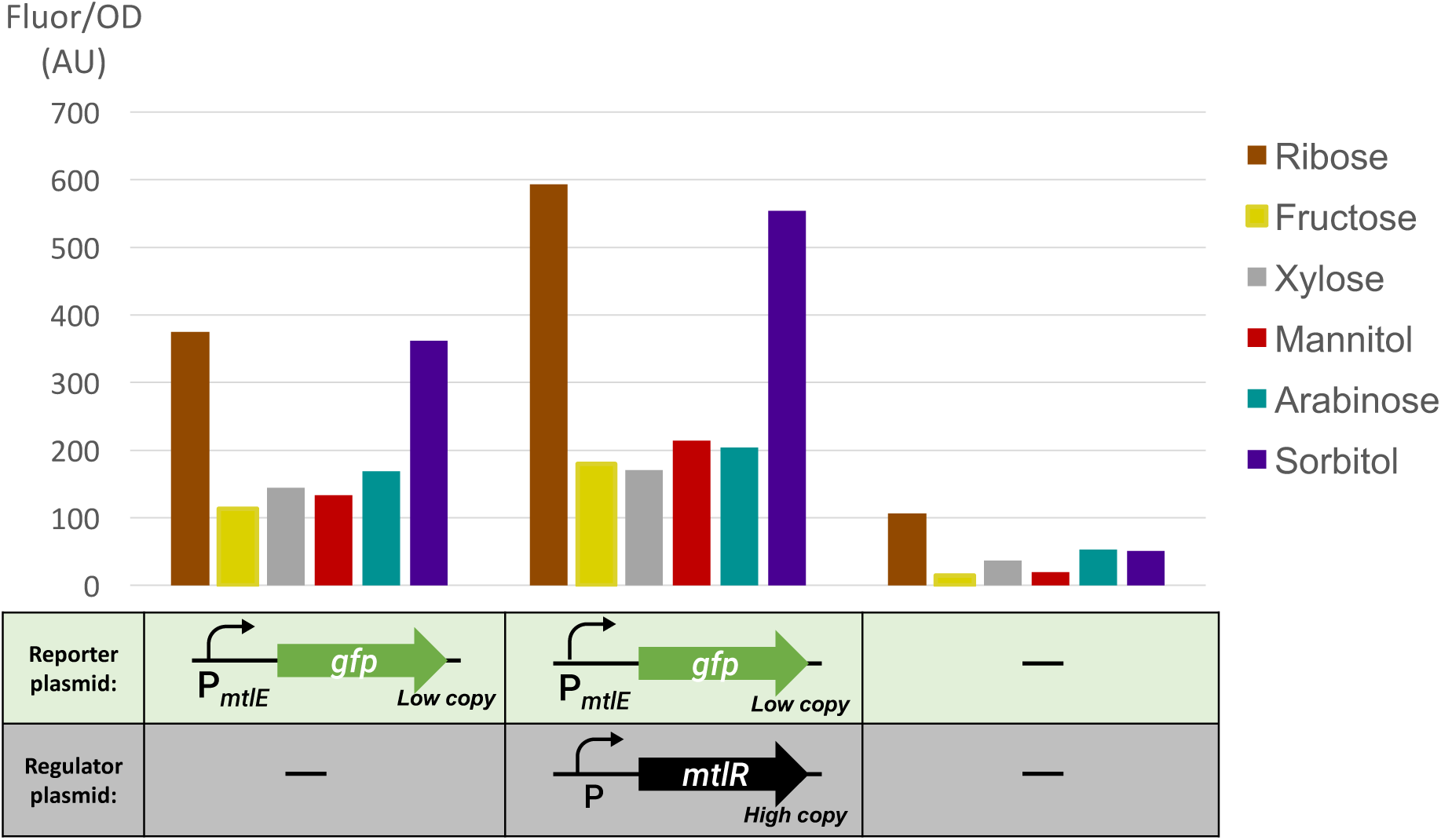
Activity od p_mtlE_/MtlR reporter circuit in E. coli. Average relative fluorescence over optical density at hour 8. Shows fluorescence levels in presence of each sugar tested (as specified). The cells were expressing either reporter plasmid BBa_K2442206 alone, both reporter and regulatory plasmid BBa_K2442202, or neither (as specified). Low copy; PSB1C3 plasmid vector. High copy; PSB3k3 plasmid vector. E. coli strain is DH5α. n=1.

### Split regulatory components of the L-arabinose operon from *E. coli* are functional and can be used for construction of L-arabinose-inducible systems

As the *P. fluorescens* MtlR regulatory system lacked specificity in *E. coli*, we aimed to utilize components of the L-arabinose operon from *E. coli* to generate a new tightly controlled xylulose-regulatory system. We chose to mutagenize the AraC protein to change its effector specificity, based on previous reports of successful engineering of AraC to respond to non-native inducers [16][17][18]. The p_*BAD*_ promoter is regulated by the AraC transcriptional regulator, which drives expression from p_*BAD*_ only in presence of L-arabinose [15]. In nature the p_*BAD*_ promoter overlaps with the *araC* coding region [20]. To allow for mutagenesis, we split *araC* from p_*BAD*_. The minimal p_*BAD*_ was designed to retain all the sites required for AraC binding. The start codon of *araC* within p_*BAD*_ has been changed from ATG→AGT (S1 Fig).

To test activity of minimal p_*BAD*_, we expressed regulatory and reporter plasmids in AraC-negative *E. coli* strain DS941 and plated the cells on LB medium. GFP fluorescence measurements demonstrated that AraC expressed from a separate plasmid can induce expression from minimal p_*BAD*_ upon binding of arabinose, as cells exhibited fluorescence only in presence of arabinose (Fig 5 and S2 File). GFP fluorescence measurements revealed that minimal p_*BAD*_ is inducible by L-arabinose 300-fold (Fig 5 and S2 File), showing improvement over previously characterized p_*BAD*_ parts. The minimal p_*BAD*_ promoter is tightly regulated by AraC expressed either from our regulatory plasmid, or from bacterial chromosome.

**Figure 5.**
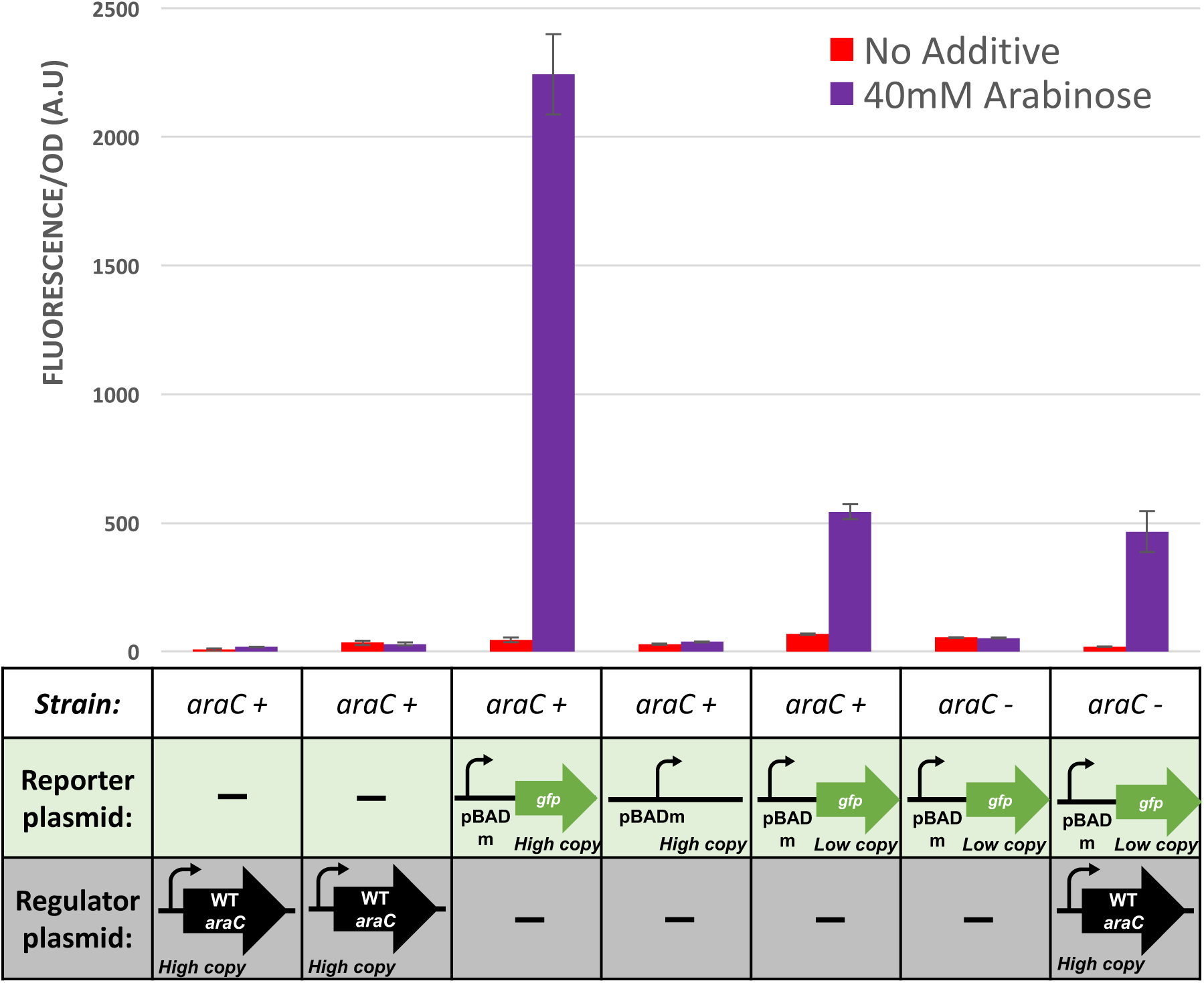
Activity of split araC and P_BAD_ fluorescence circuits in E. coli. Average relative fluorescence over optical density at hour 8. Shows fluorescence levels under no additive and 40mM Arabinose. E. coli strains which the plasmids were transformed into are DS941(araC-) or DH5α (araC+). Cells were expressing either regulatory construct BBa_K2442104, reporter construct BBa_K2442102, both, or neither (as specified). Low copy; PSB1C3 plasmid vector. High copy; PSB3k3 plasmid vector. Error bars are standard deviation; n=3

### Site-directed mutagenesis of the AraC protein provides a tool to develop biosensors responsive to non-native molecules

We performed site-directed saturation mutagenesis of the *araC* gene, targeting four amino acids within the L-arabinose binding pocket of AraC. The protocol successfully generated a mutant library of ∼24,000 AraC variants. Sequencing confirmed that NNS mutations were introduced at correct codon positions (residues 8, 24, 80 and 82; Fig 6). *E. coli* DS941 transformed with the mutant library and reporter plasmids were screened for GFP fluorescence in presence of 3 different inducer molecules that were available to us, to identify colonies with altered AraC/p_*BAD*_ activity. Due to prohibitive cost, we were unable to test xylulose. Nevertheless, 4 colonies displaying altered expression patterns were identified and subsequently characterized (Fig 7 and S2 File). Two AraC mutants constitutively activated p_*BAD*_, without the requirement for L-arabinose. The other two had lost the ability to respond to arabinose. Potentially, the latter two variants could be responsive to a yet unidentified compound.

**Figure 6.**
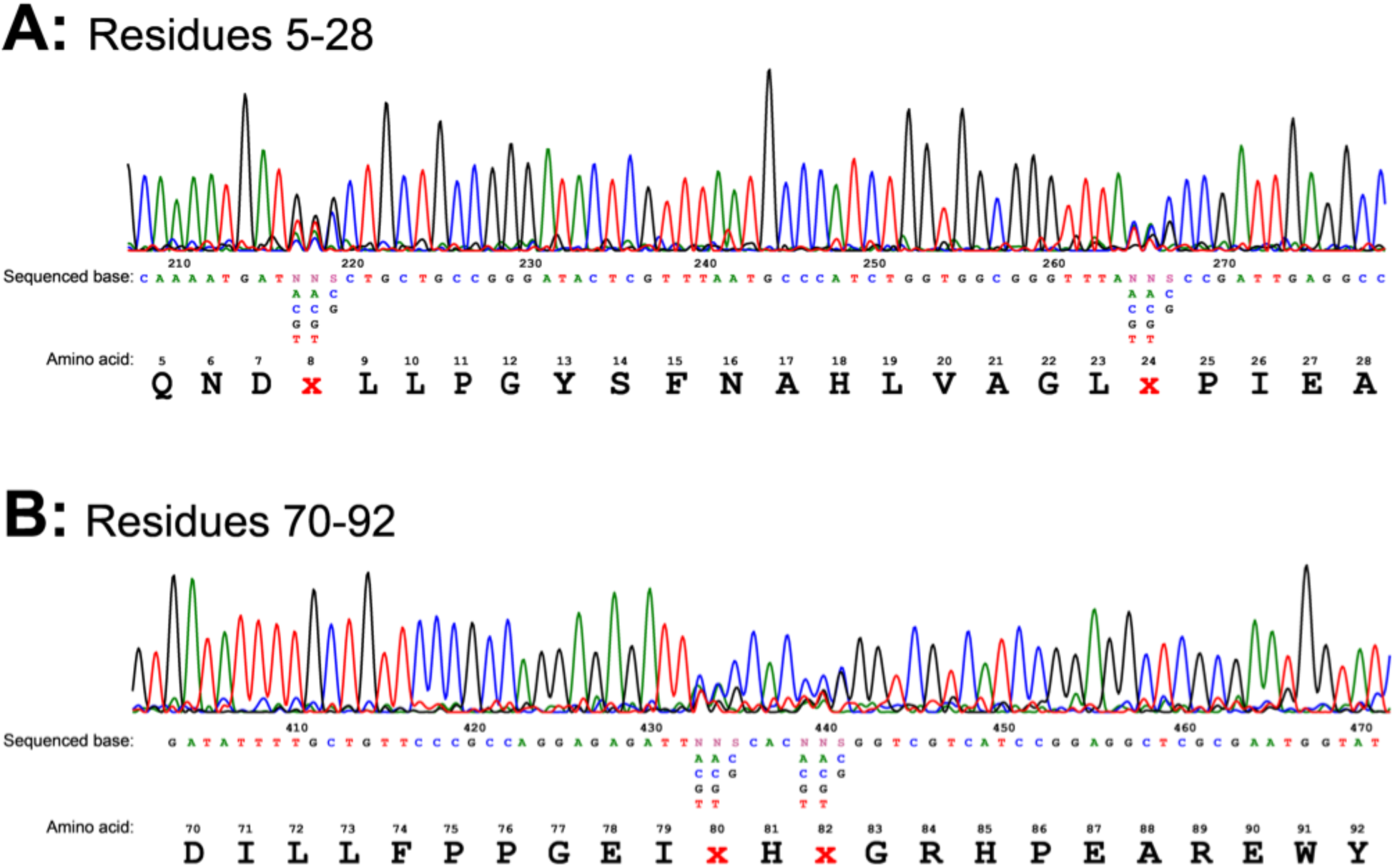
Sequencing trace from mutant library MiniPrep. Figure shows NNS mutations introduced at residues 8, 24, 80 and 82. N; any base. S; strong base (C or G).

**Figure 7.**
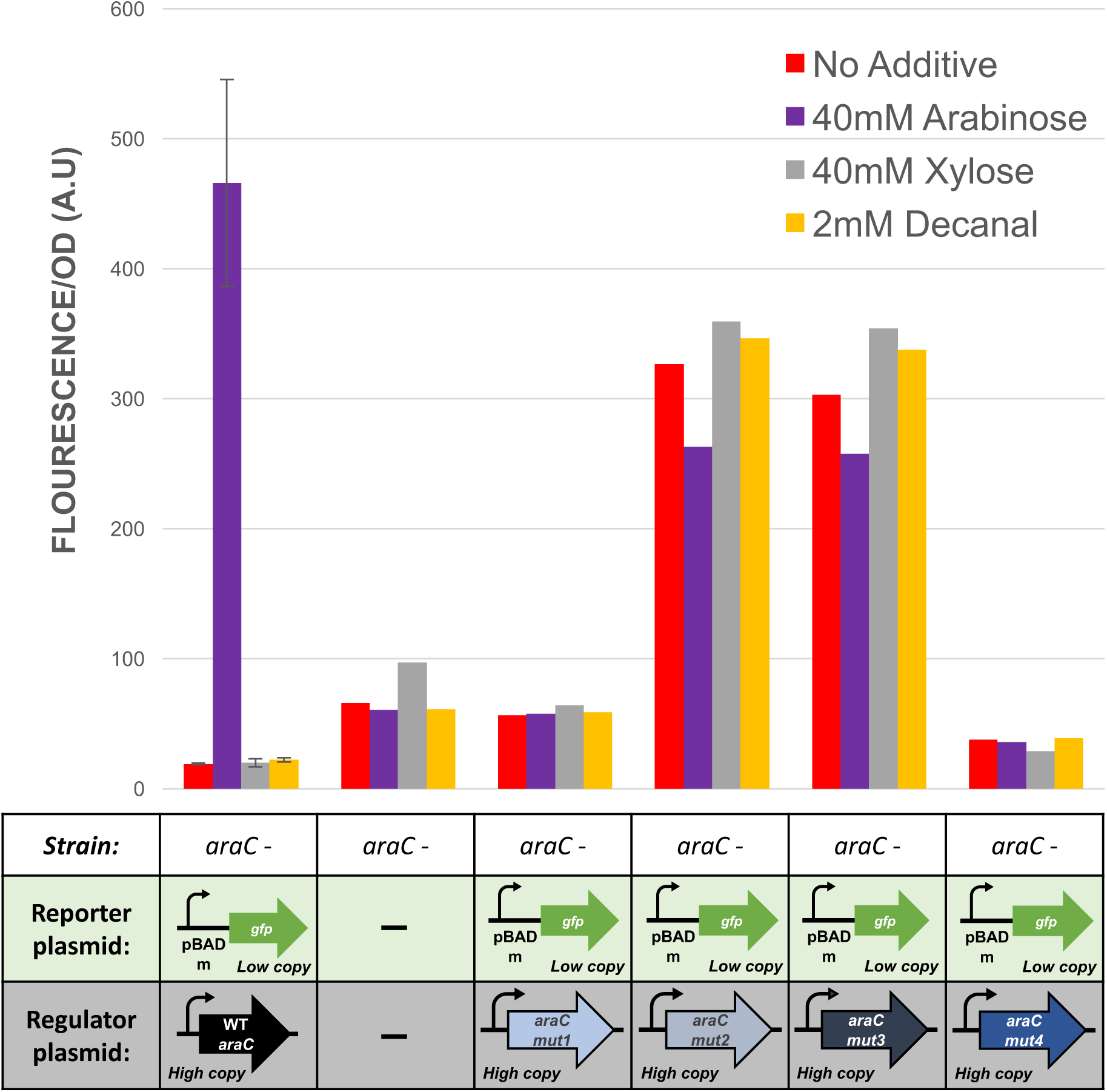
Activity of AraC mutants compared to WT AraC. Average relative fluorescence over optical density at hour 8. Shows fluorescence levels under no additive, 40mM Arabinose, 40mM xylose and 2mM decanal. E. coli strain which the plasmids were transformed into is DS941 (araC-). Cells were expressing the reporter plasmid and regulatory plasmid with wild type (WT) or mutant (mut1-mut4) variant of AraC (as specified). Empty cells served as control. Low copy; PSB1C3 plasmid vector. High copy; PSB3k3 plasmid vector. For WT AraC (lane 1) error bars are standard deviation; n=3. For all other samples (lanes 2-6): n=1.

The 4 mutant plasmids were sequenced. Wild-type AraC possesses proline at residue 8, threonine at residue 24, histidine at 80 and tyrosine at 82. Mutant 1 (BBa_K2442105) has the following mutations: proline-8→glycine (P8G), histidine-80→proline (H80P) and tyrosine-82→tryptophan (Y82W). Mutant 2 (BBa_K2442106) and 3 (BBa_K2442107) both had the same mutations as follows: proline-8→serine (P8S), threonine-24→aspartic acid (T24D), histidine-80→isoleucine (H80I) and tyrosine-82→alanine (Y82A). Sequencing of mutant 4 (BBa_K2442108) revealed a deletion within the *araC* coding region, which explains lack of AraC activity in the corresponding colonies.

## Discussion

Due to lack of cheap and rapid detection methods for *C. jejuni*, we designed a biosensor responsive to a biomarker specific to the pathogen-xylulose. In our study we characterized activity of a mannitol-responsive regulator MtlR in *E. coli*. Liu *et al*. (2015) [14] previously reported that mannitol and xylulose act as direct inducers of MtlR to activate transcription from the p_*mtlE*_ promoter. Our results suggest that this system is functional when expressed in *E. coli*, with MtlR activating transcription from p_*mtlE*_. However, reporter expression was also induced independently of MtlR. This was achieved in presence of a variety of sugars structurally similar to mannitol and xylulose, including arabinose, fructose, xylose, ribose and sorbitol. Contradictory to how the system works in *P. fluorescens* [14], the strongest reporter expression was achieved in presence of sorbitol and ribose, rather than mannitol. Although the p_*mtlE*_ was previously found to be activated by sorbitol independently of MtlR [14], no response to ribose has yet been observed. We suggest that other, yet unidentified proteins naturally found in *E. coli* act as transcriptional activators of p_*mtlE*_.

Although our results show an increase in fluorescence upon expression of the p_*mtlE*_*/*MtlR components in presence of sugars, it is important to note that our testing lacked a no-inducer control. As reporter expression from p_*mtlE*_ was observed in absence of MtlR expression, it may be that this system is constitutively activated independently of any inducer sugar. Inclusion of such control would allow for better understanding of the system’s activity in *E. coli*. Moreover, due to time constraints the experiment was only performed on only one occasion, which could impact reliability of the results. This would be improved in future experiments by increasing the number of repetitions and samples tested, to determine statistical significance of the results obtained.

As we didn’t identify a natural regulatory system responsive specifically to xylulose, we performed mutagenesis of the L-arabinose-responsive protein AraC as an alternative route to detect our target sugar. To achieve this, we split the *araC* coding region from the overlapping sequence of the p_*BAD*_ promoter. Our results suggest that the modified p_*BAD*_ is functional and can be activated by AraC expressed from a separate construct. Most importantly, we demonstrated successful mutagenesis protocol of the AraC protein to generate variants with altered effector substrate specificity. However, due to time constraints, we were only able to screen 200 mutant colonies out of 24,000 colonies obtained. This is an extremely low number considering our protocol was expected to generate 160,000 mutant AraC variants with different amino acid combinations at the 4 randomised residue positions. Moreover, limited resources enabled us to test for response to a narrow range of inducer molecules. In the future, optimization of the transformation protocol to obtain enough colonies to cover the entire library, along with a larger scale mutant screening would potentially allow identification of an AraC variant responsive to xylulose. Such mutant could be utilized in a xylulose-sensing component in a *C. jejuni* biosensor. Moreover, a catalogue of AraC mutants could be utilized as a component of a biosensor toolbox, allowing for generation of genetic circuits regulated by small molecules of choice.

Although we aimed to generate a xylulose-regulated system, we were unable to test our constructs in presence of xylulose due to its prohibitive cost. Nevertheless, the sugar would be required to screen for AraC mutants responsive to xylulose. We found that xylulose isomerase enzyme is capable of converting the cheaper sugar, xylose, into xylulose [21][22]. We have considered development of a suitable expression plasmid, which in the future could be used to overexpress and purify the enzyme for production of xylulose in subsequent experiments (for full description see http://2017.igem.org/Team:Glasgow/XyluloseBiosynthesis).

Although xylulose is rarely found in bacterial capsules [8], potential contamination of the tested area by xylulose from other sources could lead to false positive results from our detector. For this reason, we designed our biosensor to detect two sensory inputs. Apart from xylulose, we identified autoinducer-2 (AI-2) as another marker for *C. jejuni* [9]. AI-2 is a secreted quorum sensing molecule. In future development of the biosensor components, the detectors for both xylulose and autoinducer-2 would form two components of an AND gate that will ensure a positive result is given only when both xylulose and autoinducer-2 are present, improving specificity and accuracy of the detector (for full description see http://2017.igem.org/Team:Glasgow/ANDGate).

To increase efficiency of the whole-cell based biosensor, we went through several design iterations to create a device that would house the engineered bacteria to make the use of the biosensor simple and easy (for full description see http://2017.igem.org/Team:Glasgow/Applied_Design; http://2017.igem.org/Team:Glasgow/Hardware).

## Conclusions

We conclude that the AraC mutagenesis protocol was successful at generating AraC variants with altered effector specificity. A larger scale mutant screen could result in identification of a xylulose-inducible AraC variant. For increased specificity, the xylulose-regulated p_*BAD*_/AraC system could be combined with an AI-2-sensing construct within an AND gate, and transformed into a host bacterium to produce a dual-input biosensor tool for rapid detection of *C. jejuni*.

## Acknowledgements

We would like to thank Dr. Steven Kane for helping with the design of the minimal p_*BAD*_ promoter. We acknowledge Dr. Paul Everest for providing information about *C. jejuni*, Dr. Richard Daniel for donating a sample of *Bacillus subtilis* for use in our project and advice on how to work with the bacterium. We also thank Ruiyang He for his work on the AND gate subproject. We are also grateful to everyone who kindly supported us on our crowdfunding page.

## Ethics Statement

N/A

## Data Availability

Yes – all data are fully available without restriction. All relevant data are within the paper and relevant Supporting information files.

