## Supplementary Materials for "Characterizing Genetic Circuit Components in *E. coli* towards a *Campylobacter jejuni* Biosensor"

### Response to Reviewers

**PreethaG:** This abstract could be made more concise by omitting specifics of your experimental approach and instead just quickly summarizing your introduction, including a 1- or 2-sentence description of the methods, summarizing your results, and briefly mentioning the implications of those results.

**Response:** As suggested, we have made the abstract more concise by omitting some of the detail as well as specifics of our experimental approach. The methods, aims and results were briefly summarized.

**lgallagher:** Hi University of Glasgow, Thanks for your submission. Please can you make the raw plate reader data available as a link or supplementary file, rather than upon request from the corresponding author? This ensures that the data is publicly available without restrictions.

**Response:** We have attached the raw plate reader data as a supplementary PDF to ensure all data is publicly available without restrictions.

**EmmaDAL:** Nit-picky point, but seeing as you have already introduced Campylobacter jejuni, you can simply write C. jejuni for the rest of the report :)

**Response:** As suggested, we have used C. jejuni for the rest of the report.

**EmmaDAL:** Is xylulose in the polysaccharide capsule of other Campylobacter species? If so, the sensor could have a much wider reach than one bacterial species. Are you specifically trying to identify C. jejuni? Maybe need to think of additional molecules, specific to C. jejuni, to sense

**Repsonse**: To our knowledge it hasn't been studied whether other Campylobacter species have xylulose contained in their capsules, but that is definitely something to consider. The only other marker of C. jejuni that we have found was autoinducer-2, which is much less specific than xylulose. It would be worth identifying additional C. jejuni - specific markers to increase the accuracy of our sensor. Thank you for your feedback!

**EmmaDAL:** Might be mistaking the wording, but did you create this figure or is it an adaptation from the groups that you mention in the previous paragraph. If you adapted the figure, it would be helpful to mention so in the figure caption. It's a very helpful figure nonetheless

**Response**: This figure was done by us in PyMOL, by ray tracing of the araC structure. We have now made it clear in the figure caption.

**aaa_2018:** I'd recommend including the protocols in the article itself (maybe as Supplementary Information) - the iGEM website might not be online forever.

**Response:** We have included all the protocols as a Supplementary PDF.

**lgallagher:** Just a small point but we'd suggest writing in full sentences throughout so that the text flows better for the reader.

**Response:** We have edited the text to remove the short statements and make it flow better.

**lgallagher:** The text in your figures is a little small, it might be worth reformatting them and resizing parts to ensure that the text and numbers are clear

**Response:** We have edited our figures so that the labels and numbers are clear to the reader.

**madsenc1**: Did you see reproducibility for this data? I don't see errors bars and it would be nice to see that this data was reproducible.

**Response:** We edited or figures to include error bars where possible. We also included information about how many repetitions were performed in the figure legends, which hopefully makes it clear to the reader about how reproducible this data is.

**lgallagher:** PLOS doesn't allow instances of data not shown, as such we'd suggest you describe data which can be referenced or remove references to data which can't be shown.

**Response:** The references to the data which can’t be shown have been removed.

**madsenc1:** Citing my previous comment above as well as for the next figure, the figures themselves are clean and easy to read which is excellent. Yet, I would be curious to see how reproducible these experiments are.

**Response:** As above, we have edited our figures and figure legends accordingly.

**lgallagher:** Hi University of Glasgow, As this sensor has been identified for the detection of bacteria associated with food poisoning, can you describe how you think it could be used in real life situations?

**Response:** Since in the paper we describe underlying science for a potential biosensor, but we haven’t yet made a sensor, at this stage it was too early for us to predict how the biosensor could be used in real life situations.

**aaa_2018:** Nice project!

**Response:** Thank you!
