## Supplementary Materials for "Characterizing Genetic Circuit Components in *E. coli* towards a *Campylobacter jejuni* Biosensor"


#### Author contributions

Conceptualization: Natalia Brzozowska, Jane Gourlay, Ailish O'Sullivan, Frazer Buchanan, Ross Hannah, Alison Stewart, Hannah Taylor, Reuben Docea, Greig McLay, Ambra Guiliano

Methodology: Natalia Brzozowska, Jane Gourlay, James Provan

Investigation: Natalia Brzozowska, Jane Gourlay, Ailish O'Sullivan, Frazer Buchanan, Ross Hannah, Alison Stewart, Hannah Taylor, Reuben Docea, Greig McLay

Writing: Natalia Brzozowska

Formal Analysis: Natalia Brzozowska, Jane Gourlay, Ailish O'Sullivan, Frazer Buchanan, James Provan

Resources: Sean Colloms, Julien Reboud

Funding Acquisition: Sean Colloms, Julien Reboud

Supervision: Sean Colloms, Julien Reboud, James Provan, Katherine Baker, Jumai Abioye

Visualization: Natalia Brzozowska, Katherine Baker, Jane Gourlay, Ailish O'Sullivan

#### Abstract


*Campylobacter jejuni* is a bacterial pathogen responsible for the majority of food poisoning in the UK. The most important transmission route is consumption of undercooked chicken or other foods cross-contaminated from raw poultry meat. Current detection systems for the pathogen are time-consuming, expensive or inaccessible for everyday users. In this article we propose a cheaper and faster system for detection of *C. jejuni* using synthetic biology. Towards designing a dual-input biosensor, we concentrated on characterising two regulatory systems that displayed a potential to act as tools for detection of xylulose – a rare sugar naturally present in the capsule of *C. jejuni*. The regulatory protein component of the mannitol operon from *Pseudomonas fluorescens,* MtlR, was previously reported to activate expression from $p_{mtlE}$ promoter upon binding of xylulose. Using a two-part reporter system we characterised this genetic circuit in *Escherichia coli.* Our findings suggest that $p_{mtlE}$ is non-specifically regulated by other proteins found in the non-native host, and is activated in presence of a variety of inducer molecules. As an alternative, we utilized components of the L-arabinose operon naturally found in *E. coli.* The AraC regulatory protein activates expression from the $p_{BAD}$ promoter upon binding L-arabinose. We performed site-specific saturation mutagenesis to generate AraC variants with altered effector specificity. As a proof of concept, we characterised a mutant AraC variant which has lost the ability to respond to L-arabinose. A larger-scale mutant screening could result in identification of a xylulose-responsive AraC variant, which in combination with an autoinducer-2 sensing construct would allow construction of a dual-input biosensor for detection of *Campylobacter jejuni*. In addition, the protocol described here can be used as part of a biosensor toolbox to produce genetic circuits for detection of small molecules of interest.

**Comments**

**PreethaG:** This abstract could be made more concise by omitting specifics of your experimental approach and instead just quickly summarizing your introduction, including a 1- or 2-sentence description of the methods, summarizing your results, and briefly mentioning the implications of those results.

### Financial Disclosure

The project was funded by New England Biolabs, Eppendorf UK, Wellcome Trust, Institution of Mechanical Engineers, University of Glasgow, Biotechnology and Biological Sciences Research Council (BBSRC), The Microbiology Society and Society for Experimental Biology (SEB). BMG LabTech UK provided a microplate reader; IDT provided 20 kb of DNA synthesis. The funders had no role in study design, data collection and analysis, decision to publish, or preparation of the manuscript.

### Competing Interests

The authors have declared that no competing interests exist.

### Ethics Statement

N/A

### Data Availability

Yes – all data are fully available without restriction. Additional data are available on our website: http://2017.igem.org/Team:Glasgow (http://2017.igem.org/Team:Glasgow). Raw plate reader data are available upon request from the corresponding author.

**Comments**

**lgallagher:** Hi University of Glasgow, Thanks for your submission. Please can you make the raw plate reader data available as a link or supplementary file, rather than upon request from the corresponding author? This ensures that the data is publicly available without restrictions.
**Natalia:** Hi, sure we will provide a link!

### Introduction

*Campylobacter jejuni* is a Gram-negative, microaerophilic, corkscrew-shaped bacteria which has been implicated as being one of the most common causes of human gastroenteritis worldwide [1][2][3]. Infection with *Campylobacter* causes common symptoms such as diarrhoea, abdominal pain, fever, headache, nausea and vomiting [3]. *C. jejuni* is most commonly found on undercooked poultry, though it has been reported in other undercooked meat and meat products, raw milk, and in untreated drinking water [3]. The high prevalence of *C. jejuni* makes it an interesting target for synthetic biology-based solutions.

Traditional methods for detection of *Campylobacter* include culture-based techniques, which are relatively cheap to carry out and require less training than other methods [4]. However, they are incredibly time and labour intensive. Therefore, in recent years, a move has been made towards use of rapid detection testing [5]. Such techniques include enzyme immunoassay and lateral flow systems, which require only one to two hours to produce a result [6]. However, use of these methods requires highly trained employees to carry them out, and so detection of *Campylobacter* in an industrial or agricultural setting would most likely have to be outsourced to a company specialising in these services. In addition, a comparison of three rapid detection systems showed a high number of false negative results, which is a drawback when considering detection of *C. jejuni* to reduce the incidence of disease outbreaks [7].

In an attempt to reduce incidences of food poisoning by *Campylobacter jejuni*, and to improve upon current methods of detection, we decided to create a biosensor that was able to detect the presence of *Campylobacter jejuni* quickly and accurately. For this purpose we aimed to create a two-part biosensor that will detect two sensory inputs. The first, xylulose, is a rare sugar found to be incorporated into the polysaccharide capsule of *Campylobacter jejuni* [8]. The presence of xylulose is not common in bacterial polysaccharide capsules [8]. Additionally, the glycosidic bonds linking xylulose were found to be extremely acid-labile [8], providing a possibility to release the molecule from the capsule and allow for whole-cell based detection.

**Comments**

**EmmaDAL:** Nit-picky point, but seeing as you have already introduced Campylobacter jejuni, you can simply write C. jejuni for the rest of the report :)
**EmmaDAL:** Is xylulose in the polysaccharide capsule of other Campylobacter species? If so, the sensor could have a much wider reach than one bacterial species. Are you specifically trying to identify C. jejuni? Maybe need to think of additional molecules, specific to C. jejuni, to sense

**Natalia:** To our knowledge it hasn't been studied whether other Campylobacter species have xylulose contained in their capsules, but that is definitely something to consider. The only other marker of C. jejuni that we have found was autoinducer-2, which is much less specific than xylulose. It would be worth identifying additional C. jejuni - specific markers to increase the accuracy of our sensor. Thank you for your feedback!

The other molecule we identified as a marker for *Campylobacter* was autoinducer-2 (AI-2) [9]. AI-2 is a secreted quorum sensing molecule. AI-2 is a significantly less specific biomarker, as many varied gram-positive and gram-negative bacterial species sense their population density and surrounding bacterial environment using this molecule [10]. On the other hand, its ubiquity meant that AI-2 quorum sensing regulatory system having been well characterised by prior iGEM teams [11][12]. For this reason we concentrated our work on characterising the less studied components that could potentially be used for detection of xylulose - a more specific *C. jejuni* biomarker.

For the detection of xylulose two possible avenues were explored. One of them exploits the mannitol operon naturally found in *Pseudomonas fluorescens* [13]. It has been previously reported that xylulose acts as a direct inducer of the regulatory protein MtlR, activating transcription from the $p_{mtlE}$ promoter [14]. To investigate its suitability for out detection system, we characterised the $p_{mtlE}$/MtlR regulatory system in *E. coli*.

We were aware that this approach could have some disadvantages. Coming from a different organism, the $p_{mtlE}$/MtlR regulatory system could show unexpected behaviour in *E. coli* cells. For this reason we aimed to construct an alternative regulatory system which exploits components of the L-arabinose operon, native to *E. coli* [15]. Several previous studies have shown that its regulatory protein, AraC, can be engineered to activate transcription from the

$p_{BAD}$ promoter in response to non-native small molecules [16][17][18]. Site-saturation mutagenesis of residues positioned within the ligand-binding pocket of AraC (Fig 1), coupled with fluorescence-based cell sorting, allowed the groups to isolate AraC variants with altered effector specificity [16]. Based on these findings we aimed to use multiple site-saturation mutagenesis and fluorescence-based screening to generate mutant AraC responsive specifically to xylulose.

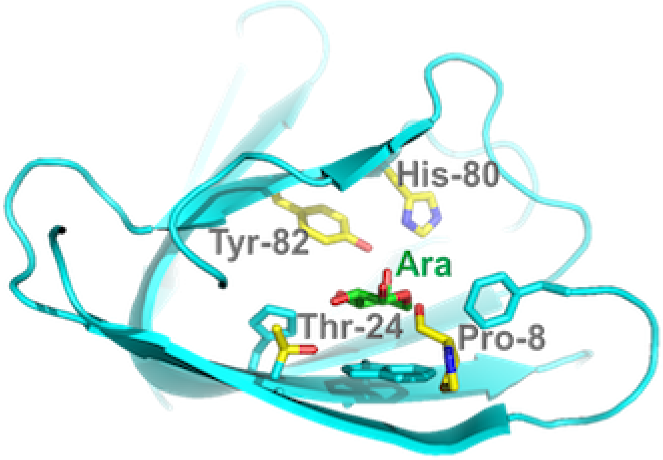


**Comments**

**EmmaDAL:** Might be mistaking the wording, but did you create this figure or is it an adaptation from the groups that you mention in the previous paragraph. If you adapted the figure, it would be helpful to mention so in the figure caption. It's a very helpful figure nonetheless
**Natalia:** This figure was done by us in PyMOL, by ray tracing of the araC structure

***Fig 1. Crystal structure of AraC binding pocket with bound L-arabinose (Ara).****The four key residues which are important for ligand binding are indicated.*

### Materials and Methods

#### Standard molecular biology techniques

During this work, a standardised set for routine laboratory techniques were used such as restriction digest, gel electrophoresis, preparation of chemically competent *E. coli*, transformation of chemically competent *E. coli*, ligation reactions, oligonucleotide annealing, and PCR. These can be found on our protocols page (http://2017.igem.org/Team:Glasgow/Protocols (http://2017.igem.org/Team:Glasgow/Protocols)). All ligation reactions described herein were transformed into DH5α commercial chemically competent *E. coli* cells, plated on L-agar containing the required antibiotics, and grown at 37°C overnight.

**Comments**

**aaa_2018:** I'd recommend including the protocols in the article itself (maybe as Supplementary Information) - the iGEM website might not be online forever.

#### Plasmid design for expression of $p_{mtlE}$/MtlR regulatory system

All plasmids were constructed using BioBrick Standard Assembly. The $p_{tet}$ promoter with a medium-strength ribosome binding site (RBS) B0032 were supplied by Integrated DNA Technologies (IDT) as oligos and ligated into a pSB1C3 plasmid backbone. *mtlR* coding sequence was supplied as a gBlock Gene Fragment by IDT and inserted behind the promoter and RBS. This produced the regulatory plasmid BBa_K2442202 (Fig 2). Sequence of the construct was verified.

The sequence of the $p_{mtlE}$ promoter with its native RBS was obtained from Liu *et al.* (2015) [14] and supplied as oligo by IDT. The reporter plasmid was constructed by ligating GFP coding sequence (BBa_E0040; obtained from iGEM Distribution Kit) downstream of the $p_{mtlE}$ promoter into pSB1C3 backbone. The resulting plasmid BBa_K2442206 is shown in Fig 2. Sequence of the final construct was verified.

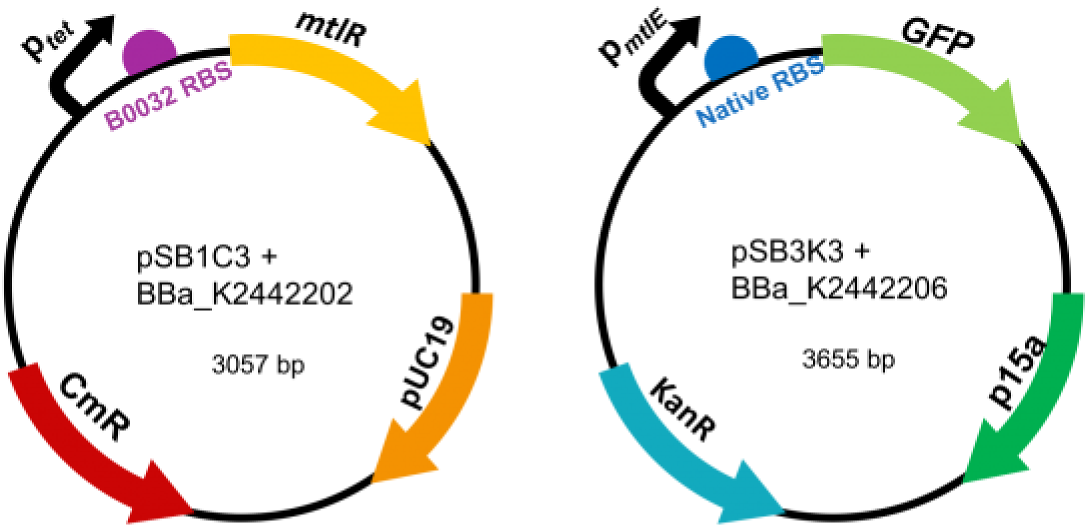


***Fig 2. Diagrams of the regulatory plasmid (left) and GFP reporter plasmid (right) for the expression of the MtlR/$p_{mtlE}$ regulatory system.***

#### Plasmid design for expression of $p_{BAD}$/AraC regulatory system

Oligos containing parts BBa_R0011 (LacI-regulated promoter) upstream of ribosome-binding site (RBS) B0032 were synthesised by IDT. Wild type *araC* sequence as described by Miyada *et al.* (1980) [19] was amplified from BBa_I0500 using primers *araC_BBPre_F* and *araC_BBSuf_R* (Table 1) to introduce BioBrick prefix and suffix to both ends of WT *araC*. The PCR product was subsequently inserted downstream of R0011 promoter and B0032 RBS in pSB1C3 backbone to generate the regulatory plasmid BB_K2442104 (Fig 3). Sequence was verified.

**Comments**

**lgallagher:** Just a small point but we'd suggest writing in full sentences throughout so that the text flows better for the reader.

Minimal $p_{BAD}$ promoter was synthesised by IDT as a gBlock. The fragment was ligated into pSB3k3, upstream of part BBa_I13500 (containing B0034 RBS and GFP). This resulted in the final reporter plasmid BBa_K2442102 (Fig 3). The sequence was verified.

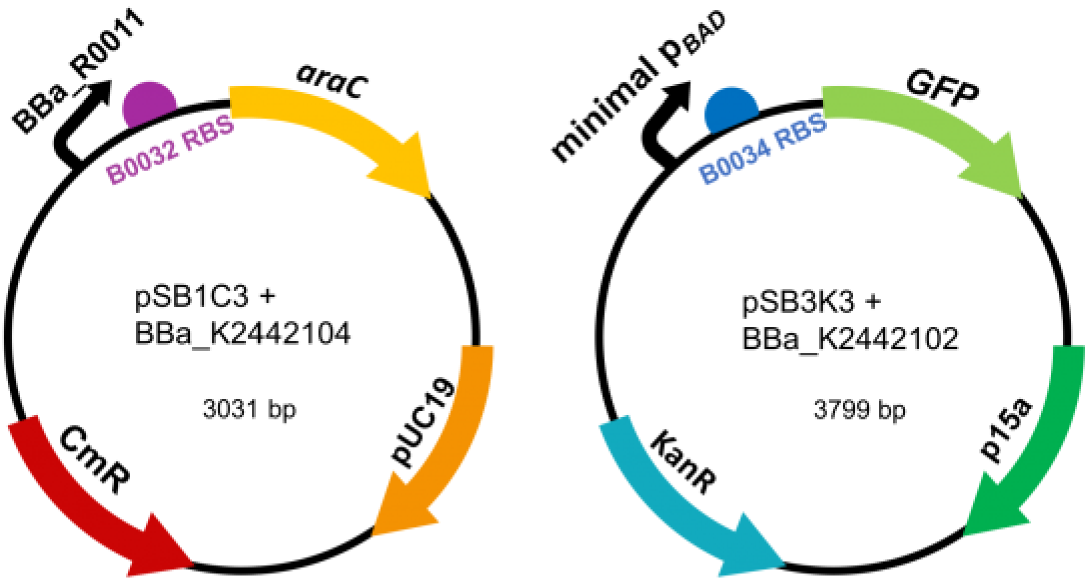


***Fig 3. Diagrams of the regulatory plasmid (left) and GFP reporter plasmid (right) for the expression of the AraC/p$_{BAD}$ regulatory system.***

#### Mutant AraC Library Construction

Primer sequences are listed in Table 1. *araC* gene was amplified from the BBa_I0500 part using primers *araC_F* and *araC_R*. The PCR product was then used in three PCR reactions with the following sets of primers: *araC_Frag1_F* and *araC_Frag1_R*; *araC_Frag2_F* and *araC_Frag2_R*; *araC_Frag3_F* and *araC_Frag3_R*. The PCR conditions were:

1. 98°C for 30s
2. 35 cycles of 98°C for 10s, 67.5°C (for F1 and F2) or 59.8°C (for F3) for 30s and 72°C for 1min
3. Final step of 72°C for 10 mins

The reactions resulted in 3 *araC* fragments (F1, F2 and F3). Primers *araC_Frag1_F*, *araC_Frag1_R* and *araC_Frag2*_R contain the degenerate sequences NNS at specific triplets so that fragments F1 and F2 are mutagenised at codon positions 8, 24, 80 and 82. PCR products were gel purified and DNA concentration of each was measured with a NanoDrop™ spectrophotometer. Equimolar aliquots (0.15 pmol each) of adjacent fragments were combined (F1+F2 and F2+F3) and PCR-assembled without primers under the following conditions:

1. 98°C for 30s
2. 15 cycles of 98°C for 10s, 60°C for 1min and 72°C for 40s
3. 72°C for 10 mins.

15 µl of each reaction product was combined and PCR-assembled without primers under the following conditions:

1. 20 cycles of 98°C for 30s
2. 72°C for 40s

Finally, primers *araC_BBPre_F* and *araC_BBSuf_R* were added and a final round of PCR was ran using the following programme:

1. 98°C for 30s
2. 30 cycles of 98°C for 10s

3. 60°C for 1 min
4. 72°C for 40s
5. 72°C for 10 min.

QIAQuick PCR purification of the product was performed according to the Qiagen protocol. The purified PCR product was ligated into pSB1C3 backbone, downstream of R0011 promoter and B0032 RBS. Ligation reaction was ethanol-precipitated, and transformed into electrocompetent DH5α by electroporation. This resulted in the *araC* mutant library. Colonies carrying the mutant library were washed off transformation plates with 1.5ml double distilled $H_2O$, ($ddH_2O$) then transferred to a 1.5ml tube. The cell pellet was centrifuged at room temperature, the supernatant was then discarded and pellet resuspended in 1.5ml $ddH_2O$. This centrifugation-resuspension step was repeated 3 times to remove agar plate debris. Plasmid DNA was extracted using the Qiagen Plasmid MiniPrep Kit with 2 of each PB buffer and PE buffer wash steps to ensure maximum extract purity. Sequencing primer VF2 was then added and sample sent for sequencing.

| Primers | Sequence |
|---|---|
| araC_F | ATGGCTGAAGCGCAAAATGAT |
| araC_R | TTATGACAACTTGACGGCTACATCA |
| araC_BBPre_F | ACGATGGAATTCGCGGCCGCTTCTAGATGGCTGAAGCGCAAAATGAT |
| araC_BBSuf_R | GGCGTACTGCAGCGGCCGCTACTAGTATTATGACAACTTGACGGCTACATCA |
| araC_Frag1_F | ATGGCTGAAGCGCAAAATGATNNSCTGCTGCCG |
| araC_Frag1_R | TAACCGTTGGCCTCAATCGGSNNTAAACCCGC |
| araC_Frag2_F | CCGATTGAGGCCAACGGTTA |
| araC_Frag2_R | CGAGCCTCCGGATGACGACCSNNGTGSNNAATCTCTCC |
| araC_Frag3_F | GGTCGTCATCCGGAGGCTCG |
| araC_Frag3_R | TTATGACAACTTGACGGCTACATCA |

***Table 1. Primer sequences used in library construction.*** *Degenerate sequences are highlighted in magenta. N, any base. S, strong base (G or C). Red, random 6bp sequence. Yellow, BioBrick prefix. Cyan, reverse compliment of BioBrick suffix.*

#### Characterisation of the $p_{mtlE}$/MtlR system activity in *E. coli*

To characterise activity of the $p_{mtlE}$/MtlR in *E. coli*, we studied the levels of GFP fluorescence using a 96 well plate in the FLUOstar Omega plate reader (BMG). Each plasmid was grown in an overnight culture of LB with appropriate antibiotics for each plasmid (chloramphenicol and kanamycin) and then diluted 1:100 with fresh LB and placed into a black bottomed 96 well plate. All readings were done in the plate reader at 37°C shaking at 200 RPM. GFP excitation and emission levels were read (excitation at 485nm and emission at 530nm) every hour for 8 hours.

### Results

##### Expression from $p_{mtlE}$ is induced by MtlR and other native regulatory proteins in *E. coli*

From the literature, we identified a regulatory system that responds to xylulose - the mannitol-inducible promoter from *Pseudomonas fluorescens* and its regulatory protein MtlR [14]. To utilise these parts in our dual-input biosensor, their activity needed to be characterised in *E. coli*. Two constructs were assembled: the regulatory plasmid with a constitutively active $p_{tet}$ promoter driving expression of the MtlR protein, and the reporter plasmid containing $p_{mtlE}$ promoter regulating expression of GFP (Fig 1).

To test whether MtlR can activate $p_{mtlE}$ in *E. coli*, we measured GFP fluorescence in *E. coli* expressing the reporter plasmid alone and both the regulatory and reporter plasmids. The experiment was done in presence of 6 structurally similar sugars (ribose, fructose, xylose, mannitol, arabinose and sorbitol) [14], to investigate substrate specificity of MtlR. Fluorescence levels were compared to basal fluorescence levels of DH5α cells, not expressing either plasmid. Cells expressing the reporter plasmid alone showed higher levels of fluorescence than empty cells, in presence of all sugars tested. This suggests that $P_{mtlE}$ is recognised and activated by other yet unidentified proteins, naturally present in *E. coli.* GFP fluorescence levels increased in cells expressing both MtlR and the reporter plasmid. This supports previous findings that MtlR functions as an activator of $p_{mtlE}$ in *E. coli.* Although we were unable to test xylulose, we found that $p_{mtlE}$ was induced in presence of a number of structurally similar sugars, showing highest response to ribose and sorbitol. We conclude that $p_{mtlE}$ promoter functions in *E. coli*, and MtlR acts as its activator. However, MtlR is not required for $p_{mtlE}$ activity when expressed in *E. coli.*

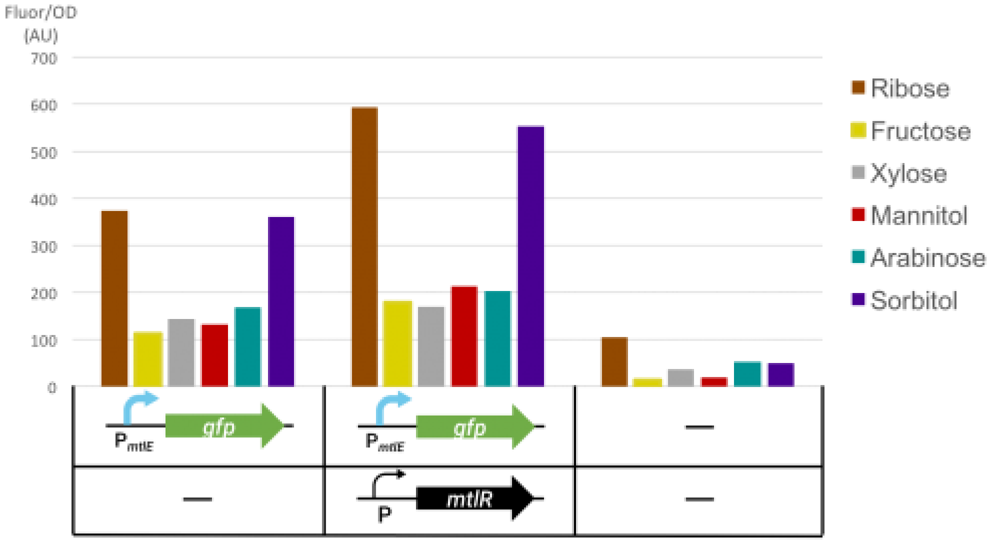


**Comments**

**lgallagher:** The text in your figures is a little small, it might be worth reformatting them and resizing parts to ensure that the text and numbers are clear

**madsenc1:** Did you see reproducibility for this data? I don't see errors bars and it would be nice to see that this data was reproducible.

***Fig 4. Activity of $p_{mtlE}$/MtlR reporter circuit in E. coli.*** *Average relative fluorescence over optical density at hour 8. Shows fluorescence levels in presence of each sugar tested (as specified). The cells were expressing either reporter plasmid BBa_K2442206 alone, both reporter and regulatory plasmid BBa_K2442202, or neither (as specified). The regulatory constructs are in pSB1C3 whilst the reporter plasmids are in PSB3K3 backbone. Cells are E. coli DH5α.*

#### Split components of the L-arabinose operon from *E. coli* are functional and can be used for construction of L-arabinose-inducible systems

As we didn't find a natural regulatory system responsive to xylulose, we aimed to utilise components of the L-arabinose operon from *E. coli* to generate our own. We chose to mutagenise the AraC protein to change its effector specificity, based on previous reports of successful engineering of AraC to respond to non-native inducers [16][17][18]. The $p_{BAD}$ promoter is regulated by the AraC transcriptional regulator, which drives expression from $p_{BAD}$ only in presence of L-arabinose [15]. In nature the $p_{BAD}$ promoter overlaps with the *araC* coding region [20]. To allow for mutagenesis, we split *araC* from $p_{BAD}$. The minimal $p_{BAD}$ was designed to retain all the sites required for AraC binding. The start codon of *araC* within $p_{BAD}$ has been changed from ATG to AGT (sequence is available at http://parts.igem.org/Part:BBa_K2442101 (http://parts.igem.org/Part:BBa_K2442101)).

To test activity of minimal $p_{BAD}$, we expressed regulatory and reporter plasmids in an *E. coli* strain DS941, which doesn't posses a functional copy of *araC* (*araC-*) ,and plated the cells on LB medium. Fluorescence imaging confirmed that AraC expressed from a separate plasmid can induce expression from minimal $p_{BAD}$ upon binding of arabinose, as cells exhibited fluorescence in presence of arabinose but not in presence of glucose (data not shown). GFP fluorescence measurements revealed that minimal $p_{BAD}$ is inducible by L-arabinose 300-fold (Fig 5), showing improvement over previously characterised $p_{BAD}$ parts. The promoter can be activated by AraC expressed either from our regulatory plasmid, or from bacterial chromosome.

**Comments**

**lgallagher:** PLOS doesn't allow instances of data not shown, as such we'd suggest you describe data which can be referenced or remove references to data which can't be shown.

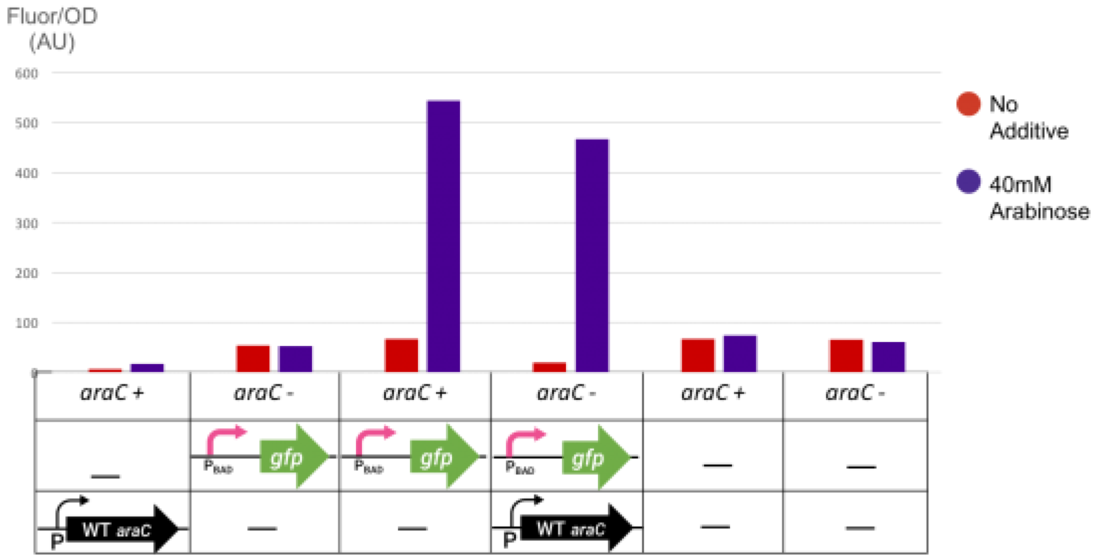


**Comments**

**madsenc1:** Citing my previous comment above as well as for the next figure, the figures themselves are clean and easy to read which is excellent. Yet, I would be curious to see how reproducible these experiments are.

***Fig 5. Activity of split p$_{BAD}$/AraC reporter circuit in E. coli.*** *Average relative fluorescence over optical density at hour 8. Shows fluorescence levels under no additive or 40mM Arabinose (as specified). The cells which the plasmids have been transformed into are E. coli DS941(araC-) or DH5α (araC+). Cells were expressing either regulatory construct BBa_K2442104 in pSB1C3, reporter construct BBa_K2442102 in pSB3k3, both, or neither (as specified).*

#### Site-directed mutagenesis of the AraC protein provides a tool to develop biosensors responsive to non-native molecules

We performed site-directed saturation mutagenesis of the *araC* gene, targeting four amino acids within the L-arabinose binding pocket of AraC (Fig 1). The protocol generated a mutant library of ~24,000 AraC variants. Sequencing confirmed that NNS mutations were introduced at correct codon positions (residues 8, 24, 80 and 82; Fig 6). *E. coli* DS941 transformed with the mutant library and reporter plasmids were screened for GFP fluorescence in presence of 3 different inducer molecules that were available to us, to identify colonies with altered AraC/p$_{BAD}$ activity. Due to prohibitive cost, we were unable to test xylulose. Nevertheless, 4 colonies displaying altered expression patterns were identified and subsequently characterised (Fig 7). Two AraC mutants constitutively activated p$_{BAD}$, without the requirement for L-arabinose. The other two had lost the ability to respond to arabinose. Potentially, the latter two variants could be responsive to a yet unidentified compound.

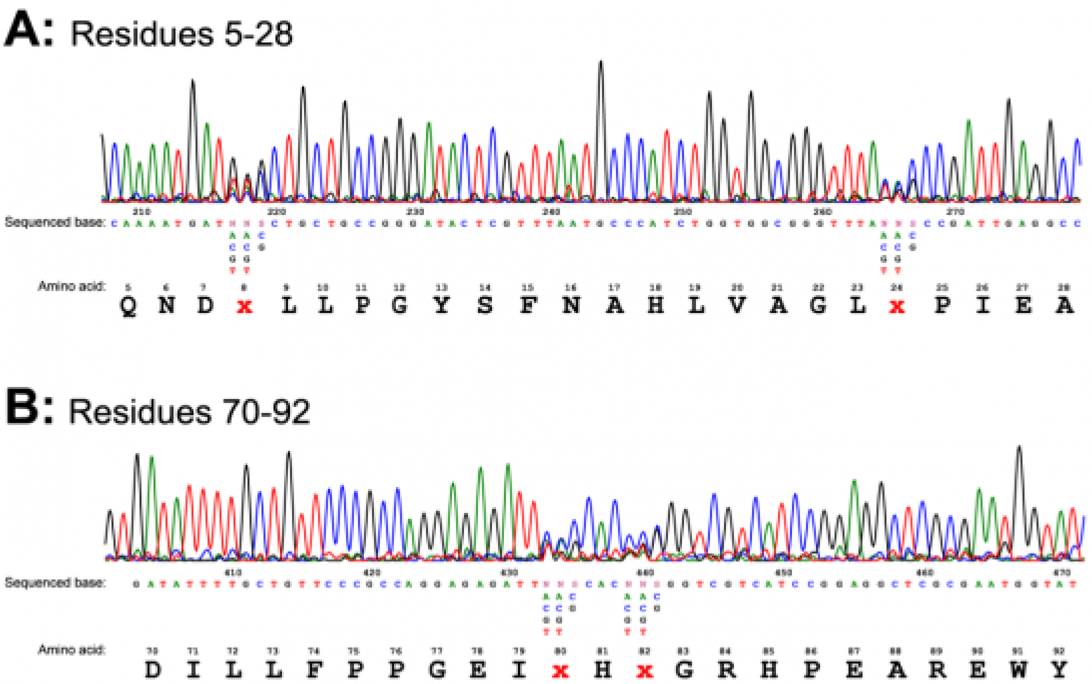


***Fig 6. Sequencing trace from mutant library MiniPrep.*** *NNS mutations at residues 8, 24, 80 and 82 can be seen.*

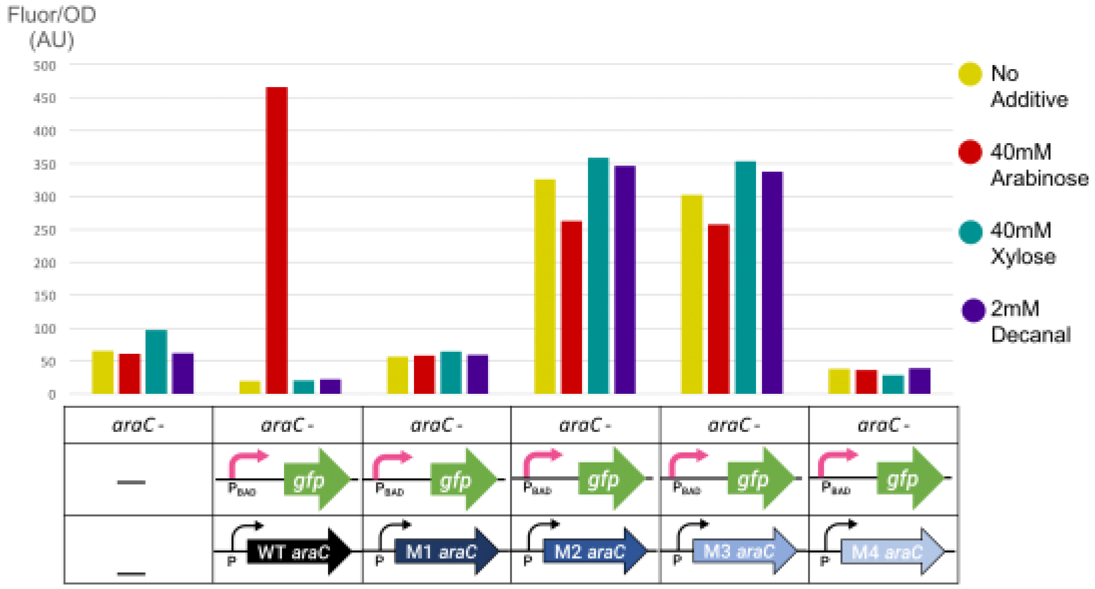


***Fig 7. Activity of mutant AraC variants in E. coli, in presence of native and non-native inducer molecules.*** *Average relative fluorescence over optical density at hour 8. Shows fluorescence levels under no additive, 40mM Arabinose, 40mM Xylose and 2mM decanal. The cells which the plasmids have been transformed into are E. coli DS941 (araC -). Cells were expressing the reporter construct BBa_K2442102 in pSB3k3 and regulatory construct with wild type (WT) or mutant (M1-M4) variants of AraC (as specified). Untransformed cells served as control.*

The 4 mutant plasmids were sequenced. AraC naturally has proline at residue 8, threonine at residue 24, histidine at 80 and tyrosine at 82. Mutant 1 (BBa_K2442105) has the following mutations: proline-8→glycine (P8G), histidine-80→proline (H80P) and tyrosine-82→tryptophan (Y82W). Mutant 2 (BBa_K2442106) and 3 (BBa_K2442107) both had the same mutations as follows: proline-8→serine (P8S), threonine-24→aspartic acid (T24D), histidine-80→isoleucine (H80I) and tyrosine-82→alanine (Y82A). Sequencing of mutant 4 (BBa_K2442108) revealed a deletion within the *araC* coding region, which explains lack of AraC activity in the corresponding colonies.

### Discussion

Due to lack of cheap and rapid detection methods for *C. jejuni,* we designed a biosensor responsive to a biomarker specific to the pathogen - xylulose. In present study we characterised activity of a mannitol-responsive regulator MtlR in *E. coli*. Liu *et al*. (2015) [14] has previously reported that in *Pseudomonas fluorescens,* an organism in which the mannitol operon is naturally found, mannitol and xylulose act as direct inducers of MtlR to activate transcription from the $p_{mtlE}$ promoter. Our results suggest that this system is functional when expressed in *E. coli*, with MtlR activating transcription from $p_{mtlE}$. However, reporter expression was also induced independently of MtlR. This was achieved in presence of a variety of sugars structurally similar to mannitol and xylulose, including arabinose, fructose, xylose, ribose and sorbitol. Contradictory to how the system works in *P. fluorescens* [14], the strongest reporter expression was achieved in presence of sorbitol and ribose, rather than mannitol. Although the $p_{mtlE}$ was previously found to be activated by sorbitol independently of MtlR [14], no response to ribose has yet been observed. We suggest that other, yet unidentified proteins naturally found in *E. coli* act as transcriptional activators of $p_{mtlE}$. Moreover, it may be that this system is constitutively on independently of any inducer sugar. Future work should probe the possibility that that unidentified cellular components could be interfering with promoter activity as this was not exhaustively characterised in this work. In addition, to determine significance of the results the number of repetitions would be increased.

As we didn’t identify a natural regulatory system responsive specifically to xylulose, we performed mutagenesis of the L-arabinose-responsive protein AraC as an alternative route to detect our target sugar. To achieve this, we split the *araC* coding region from the overlapping sequence of the $p_{BAD}$ promoter. We show that the modified $p_{BAD}$ is functional and can be activated by AraC expressed from a separate construct. Most importantly, we demonstrated successful mutagenesis of the AraC protein to generate variants with altered substrate specificity. In particular, one mutant has lost the ability to respond to L-arabinose, and may potentially be responsive to a yet unidentified, non-native molecule. Larger scale mutant screening would potentially allow identification of an AraC variant responsive to xylulose. Such mutant could be utilised in a xylulose-sensing component as part of a *C. jejuni* biosensor. Moreover, a catalogue of AraC mutants could serve in a biosensor toolbox, allowing for generation of genetic circuits regulated by small molecules of choice.

Although we aimed to generate a xylulose-regulated system, we were unable to test our constructs in presence of xylulose due to its prohibitive costs. Nevertheless, the sugar would be required to screen for AraC mutants responsive to xylulose. We found that xylulose isomerase enzyme is capable of converting the cheaper sugar, xylose, into xylulose [21][22]. We have considered development of a suitable expression plasmid, which could be used for overexpression and purification of the enzyme. This would allow synthesise of xylulose to be used in subsequent experiments (for full description see http://2017.igem.org/Team:Glas... (http://2017.igem.org/Team:Glasgow/XyluloseBiosynthesis)).

Although xylulose is rarely found in bacterial capsules [8], potential contamination of the tested area by xylulose from other sources could lead to false positive results from our detector. To improve accuracy of the reading, we designed our biosensor to detect two sensory inputs. Apart from xylulose, we identified AI-2 as another marker for *Campylobacter* [9]. AI-2 is a secreted quorum sensing molecule. In future development of the biosensor, the detectors for both xylulose and AI-2 would form two components of an AND gate that will ensure a positive result is given only when both xylulose and AI-2 are present (for full description see http://2017.igem.org/Team:Glas... (http://2017.igem.org/Team:Glasgow/ANDGate)).

To further increase efficiency of the whole-cell based biosensor, we went through several design iterations to create a self-contained device that would house the engineered bacteria to make the use of the biosensor simple and easy (for full description see http://2017.igem.org/Team:Glasgow/Applied_Design (http://2017.igem.org/Team:Glasgow/Applied_Design) and http://2017.igem.org/Team:Glasgow/Hardware (http://2017.igem.org/Team:Glasgow/Hardware)).

### Conclusions

We conclude that the AraC mutagenesis protocol was successful at generating AraC variants with altered effector specificity. A larger scale mutant screening could result in identification of a xylulose-inducible AraC variant. For increased specificity, the xylulose-regulated $p_{BAD}$/AraC system could be combined with an AI-2-sensing construct within an AND gate, and transformed into a host bacterium to produce a dual-input biosensor tool for rapid detection of *Campylobacter jejuni*.

### Acknowledgements

We would like to thank Dr. Steven Kane for helping with the design of the minimal $p_{BAD}$ promoter. We acknowledge Dr. Paul Everest for providing information about *Campylobacter jejuni and* Dr. Richard Daniel for donating a sample of *Bacillus subtilis* for use in our project and advice on how to work with the bacterium. We also thank Ruiyang He for his work on the AND gate design.

**Comments**

**lgallagher:** Hi University of Glasgow, As this sensor has been identified for the detection of bacteria associated with food poisoning, can you describe how you think it could be used in real life situations?

**Natalia:** We will touch on that in the discussion, thank you for your feedback!

**aaa_2018:** Nice project!

**Natalia:** Thank you!
