## Supplementary Materials for "Characterizing Genetic Circuit Components in *E. coli* towards a *Campylobacter jejuni* Biosensor"

Outlined below are the techniques we employed in our work.

### Preparation of CaCl2 competent cells

1. Dilute 500µl of overnight liquid culture into 20ml of broth with any necessary antibiotics to select for any plasmids already transformed into the cells.
2. Incubate at 37ºC, shaking at 225rpm for 105 minutes.
3. Spin down for 2 minutes at 7000G at 4ºC.
4. Discard supernatant, resuspend pellet in 10ml of 50 mM $CaCl_2$, keep on ice.
5. Repeat centrifugation for 2 minutes at 7000G at 4ºC.
6. Discard supernatant and the resuspend pellet in 1ml of 50 mM $CaCl_2$, keep on ice.
7. $CaCl_2$ competent cells can be kept on ice in the fridge overnight.

### Transformation of CaCl2 competent cells

1. Add 1µl of plasmid DNA to 100µl of competent cells.
2. Heat shock at 42°C for 45 seconds.
3. Add 200µl of L-broth of the sample.
4. Keep on ice for 2 minutes.
5. Incubate the cells at 37°C. Incubation time varies depending on which antibiotic resistance the plasmid holds:
    - Chloramphenicol: 90 minutes
    - Kanamycin: 60 minutes
    - Ampicillin: 30 minutes

### Restriction digests

1. For a 20µl reaction add:
    - 2µl buffer
    - 4µl Miniprep (or 8µl G-Block) dependent on concentration of Miniprep
    - Make up to 20µl with ddH20
2. Add 10-20 Units restriction enzyme(s)
3. Vortex briefly.
4. Incubate at 37°C for at least 60 minutes.
5. Heat inactivate restriction digests where appropriate.

### MiniPrep

We used the protocol from a standard Qiagen QIAprep Spin Miniprep Kit.

1. Pipette 1mL of bacterial overnight culture into a microcentrifuge tube
2. Centrifuge at 13,000 rpm for 1 min
3. Discard the supernatant
4. Repeat steps 1-3 two more times
5. Add 250µl of P1 buffer and pipette up and down to resuspend the pellet
6. Add 250µl of P2 buffer and invert Note: don't allow this lysis reaction to proceed for more than 5 minutes
7. Add 350µl of N3 buffer to neutralise the reaction and invert
8. Centrifuge at 13,000 rpm for 10 minutes
9. Transfer 800µl of supernatant into a column
10. Centrifuge for 1 minute and discard flow-through
11. Add 500µl of PB buffer and centrifuge for 1 minute

12. Discard flow-through
13. Add 750μl of PE buffer and centrifuge for 1 minute
14. Discard flow-through
15. Centrifuge again for 1 minute to get rid of any residual buffer
16. Transfer the column to a microcentrifuge tube
17. Add 50μl of EB buffer and let it stand for 1 minute
18. Centrifuge for 1 minute

### Making a gel

To make a 1.5L buffer:

1. Measure 30mL of 50x TAE buffer
2. Transfer to a 2L measuring cylinder
3. Fill up to 1.5L with distilled water
4. To make a 1% agarose gel:
5. Weigh out 1g of agarose powder
6. Transfer to a microwave bottle
7. Pour 100mL of buffer into the bottle
8. Microwave until clear
9. Cool down to 55ºC before pouring the gel

### Gel electrophoresis

To run the gel:

1. Place the gel in the tank and cover it with the buffer
2. Load 5μl of DNA ladder into the 2nd well (leave the first one empty)
3. Add 5μl of loading dye (30% glycerol, 1% bromophenol blue, 0.5% sodium dodecyl sulphate, diluted in TE buffer) into each restriction digest and pipette up and down a few times
4. Load 25μl of each restriction digest into a separate well
5. Run gel at 100V ( ~100 mA) for approximately 1 hour

To stain the gel:

- SYBR Safe: stain in (concentration) for 40 minutes, destain for 40 minutes, image under UV light
- Azure A (for gel extraction): stain in 0.04%/20% ethanol for 15 minutes, destain for 15 minutes (multiple rounds of destaining may be required), image under visible light

### Gel extraction using Qiagen kit

1. Weigh an empty eppendorf and record the weight
2. Cut out a the band from the gel and place it in the eppendorf
3. Weigh the eppendorf again and calculate the weight difference
4. Add 3 volumes of Buffer QG to 1 volume of gel
5. Incubate at 50ºC for 10 minutes until the agarose has dissolved. Vortex every 2-3 minutes
6. Add 1 volume of isopropanol and mix
7. Transfer the sample from the eppendorf to a QIAprep spin column in a 2mL collection tube
8. Centrifuge for 1 minute at 13,000rpm and discard flowthrough
9. Add 500μl of Buffer QG to column
10. Centrifuge for 1 minute at 13,000rpm and discard flowthrough

11. Add 750µl of Buffer PE and let it stand for 2-5 minutes
12. Centrifuge for 1 minute at 13,000rpm and discard flowthrough. Repeat this step
13. Transfer the column to a new eppendorf and discard the collection tube
14. Add 30µl of Buffer EB to the sample and let it stand for 1 minute
15. Centrifuge for 1 minute at 13,000rpm and keep discard the column

### Ligation

1. For a 10µl ligation reaction add (these volumes can be changed depending on the concentration of the gel extracted DNA):
    - 3µl of vector
    - 5µl of insert
    - 1µl of 10x ligation buffer
    - 0.5µl of ddH2O
    - 0.5µl of ligase
2. Vortex

1. For ligating oligo and vector add:
    - 1µl of oligo (annealed and diluted to 0.1 µM)
    - 5µl of vector
    - 1µl of 10x ligation buffer
    - 3µl of ddH2O
    - 0.5µl of ligase
2. Vortex

3. Ligation reactions were left for a minimum of 1 hour or ideally overnight at room temperature or in a temperature-stable room at 16ºC to ensure maximum ligation efficiency.

### Annealing oligos

1. Dissolve dried oligo in TE buffer to make a 100µM solution and leave for 10 minutes
2. To make a 100µl reaction add:
    - 10µl of top strand 100µM oligo
    - 10µl of bottom strand 100µM oligo
    - 80µl of TE buffer
3. Heat at 86ºC in a water bath for 5 minutes using a float
4. Scoop out some water and the float from the water bath using a large beaker
5. Let the water cool down slowly to approximately 30ºC
6. Dilute annealed oligos 1/100 to 0.1 µM

### Ethanol precipitation of ligation

1. To ligation reaction add 1/10 volume of 5M NaCl
2. Add 2 volumes of 100% ethanol
3. Vortex, place at -70$_{o}$C for 20 mins
4. Spin in chilled micro-centrifuge for 45 mins at 3000rpm
5. Pour off supernatant, spin 1 min, pipette off remainder
6. Add 1 volume 70% ethanol

7. Spin again for 30 mins, discard supernatant
8. Air dry
9. Resuspend in 10 ul of nuclease free H20

### Transformation of Library competent DH5a cells

Protocol sourced from Thermofisher scientific

1. Thaw cells on competent wet ice. Place required number of 17 times 100nm polypropylene tubes on ice
2. Gently mix cells, then aliquot 100 ul of competent cells into chilled tubes
3. For DNA from ligation reactions, dilute the reactions 5-fold in 10mM TE buffer. Add 1ul of the dilution to the cells (1-10 ng DNA), moving the pipette through the cells whilst dispensing. Gently tap to mix
4. Incubate on ice for 30 mins
5. Heat shock cells for 45 seconds in a 42oC water bath; do not shake
6. Place on ice for 2 mins
7. Add 0.9ml of room temp S.O.C Medium
8. Shake at 225rpm for 1 hour
9. Dilute the reactions as necessary and spread 100-200 ul of this dilution into LB plates with required antibiotic.
10. Incubate overnight

### Site-directed mutagenesis by PCR

Primer sequences are listed in Table 1. *araC* gene was amplified from the BBa_I0500 part using primers *araC_F* and *araC_R*. The PCR product was then used in three PCR reactions with the following sets of primers: *araC_Frag1_F* and *araC_Frag1_R*; *araC_Frag2_F* and *araC_Frag2_R*; *araC_Frag3_F* and *araC_Frag3_R*. The PCR conditions were:

1. 98°C for 30s
2. 35 cycles of 98°C for 10s, 67.5°C (for F1 and F2) or 59.8°C (for F3) for 30s and 72°C for 1min
3. Final step of 72°C for 10 mins

The reactions resulted in 3 *araC* fragments (F1, F2 and F3). Primers *araC_Frag1_F*, *araC_Frag1_R* and *araC_Frag2*_R contain the degenerate sequences NNS at specific triplets so that fragments F1 and F2 are mutagenized at codon positions 8, 24, 80 and 82. PCR products were gel purified and DNA concentration of each was measured with a NanoDrop™ spectrophotometer. Equimolar aliquots (0.15pmol each) of adjacent fragments were combined (F1+F2 and F2+F3) and PCR-assembled without primers under the following conditions:

1. 98°C for 30s
2. 15 cycles of 98°C for 10s, 60°C for 1min and 72°C for 40s
3. 72°C for 10 mins.

15 µl of each reaction product was combined and PCR-assembled without primers under the following conditions:

1. 20 cycles of 98°C for 30s
2. 72°C for 40s

Finally, primers *araC_BBPre_F* and *araC_BBSuf_R* were added and a final round of

PCR was ran using the following programme:

1. 98°C for 30s
2. 30 cycles of 98°C for 10s
3. 60°C for 1 min
4. 72°C for 40s
5. 72°C for 10 min.

| Primers | Sequence |
|---|---|
| araC_F | ATGGCTGAAGCGCAAAATGAT |
| araC_R | TTATGACAACTTGACGGCTACATCA |
| araC_BBPre_F | ACGATGGAATTCGCGGCCGCTTCTAGATGGCTGAAGCGCAAAATGAT |
| araC_BBSuf_R | GGCGTACTGCAGCGGCCGCTACTAGTATTATGACAACTTGACGGCTACATCA |
| araC_Frag1_F | ATGGCTGAAGCGCAAAATGATNNSCTGCTGCCG |
| araC_Frag1_R | TAACCGTTGGCCTCAATCGGSNNTAAACCCGC |
| araC_Frag2_F | CCGATTGAGGCCAACGGTTA |
| araC_Frag2_R | CGAGCCTCCGGATGACGACCSNNGTGSNNAATCTCTCC |
| araC_Frag3_F | GGTCGTCATCCGGAGGCTCG |
| araC_Frag3_R | TTATGACAACTTGACGGCTACATCA |

***Table 1.*** *Primer sequences used in library construction. Degenerate sequences are highlighted in magenta. N, any base. S, strong base (G or C). Red, random 6bp sequence. Yellow, BioBrick prefix. Cyan, reverse compliment of BioBrick suffix.*

### QIAQuick PCR purification

1. Add 5 volumes of Buffer PB to 1 volume of the PCR sample and mix.
2. Place a QIAquick spin column in a provided 2 ml collection tube.
3. To bind DNA, apply the sample to the QIAquick column and centrifuge for 30–60 s.
4. Discard flow-through. Place the QIAquick column back into the same tube.
5. To wash, add 0.75 ml Buffer PE to the QIAquick column and centrifuge for 30–60 s.
6. Discard flow-through and place the QIAquick column back in the same tube. Centrifuge the column for an additional 1 min.
7. Place QIAquick column in a clean 1.5 ml microcentrifuge tube.
8. To elute DNA, add 50 µl Buffer EB (10 mM Tris·Cl, pH 8.5) or water (pH 7.0–8.5) to the center of the QIAquick membrane and centrifuge the column for 1 min. Alternatively, for increased DNA concentration, add 30 µl elution buffer to the center of the QIAquick membrane, let the column stand for 1 min, and then centrifuge.
