## Supplementary Materials for "Characterizing Genetic Circuit Components in *E. coli* towards a *Campylobacter jejuni* Biosensor"

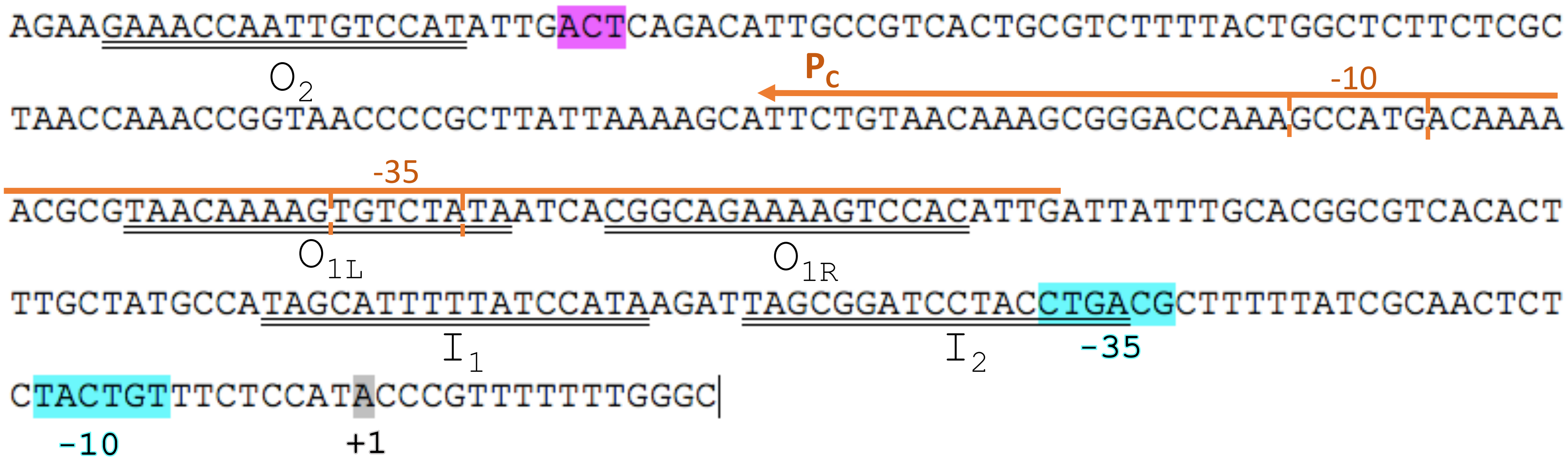


**S1 Fig. Annotated sequence of minimal $P_{BAD}$.** AraC potential start codon mutation ATG→AGT position is highlighted in magenta (in reverse compliment). AraC-binding half sites are underlined and annotated. -35 and -10 elements of the $P_{BAD}$ promoter are highlighted in cyan. AraC transcription start site (+1) highlighted in grey. $P_C$ promoter and its direction is indicated by an orange arrow. -35 and -10 elements of the $P_C$ promoter are indicated.
